# Transdiagnostic connectome-based predictive modeling reveals where circuits related to self-reported clinical symptoms impinge upon brain networks supporting cognition

**DOI:** 10.1101/2025.10.02.680080

**Authors:** Alexander J Simon, Anja Samardzija, Santino Iannone, Flor Parra Rodriguez, Saloni Mehta, Fuyuze Tokoglu, Maolin Qiu, Jagriti Arora, Kira Y Tang, Anna Q Flanagan, Rachel Katz, Gerard Sanacora, Scott W. Woods, Vinod H. Srihari, Xilin Shen, Dustin Scheinost, R. Todd Constable

## Abstract

A key assumption of the NIMH’s RDoC framework is that disordered circuits in the brain should manifest in observable behaviors, including psychiatric symptomatology and cognitive deficits. However, how disordered circuitry impacts multiple behaviors remains poorly understood. Connectome-based predictive modeling (CPM) applied to functional MRI connectivity data can identify networks associated with specific behavioral measures across individuals. Prediction strength reflects how closely a measure relates to network connectivity, while derived networks provide evidence of where an individual’s disordered circuits are located. Using CPM, we predicted a broad range of self-reported clinical and objective cognitive measures in a large, transdiagnostic sample with extensive fMRI data (n = 317). Prediction performance varied substantially across instruments, with objective cognitive tests yielding stronger models than self-reported clinical measures (*p* < 0.001). To test whether circuits underlying cognitive deficits related to symptomatology reside in regions where networks overlap, we examined the prediction strength of these sparsely shared circuits. Their connectivity strongly predicted cognitive performance and were primarily localized within the frontoparietal network and between the frontoparietal and default mode networks. These findings reveal how much various behavioral measures reflect brain networks and how circuits within the shared network space contribute to cognitive deficits associated with symptomatology.

## Introduction

Identifying neurobiological correlates of real-world behavioral measures can teach us about how neural dysfunction manifests as clinical symptoms and cognitive deficits. Such information can improve precision psychiatry by increasing our understanding of the etiology of disorders, refining diagnostic and treatment strategies, and guiding novel therapeutic development. The recognition that psychiatric symptoms and cognitive deficits stem from deviations from normal neural functioning that span diagnostic categories led to the conceptualization of the NIMH’s RDoC initiative^1,2^. In accordance with this framework, studies have utilized functional MRI (fMRI) to link high-dimensional patterns of functional connectivity distributed across the brain to a wide range of continuous behavioral measures^3–9^.

Connectome-based predictive modeling (CPM) is a well-established approach for linking behavioral measures to underlying brain networks inferred from fMRI connectivity data in a predictive modeling framework^3,10^. The cross-validation employed in building the predictive models assures that the identified brain-behavior associations are generalizable^11,12^. A large body of work has demonstrated CPM’s ability to relate functional networks to numerous cognitive and clinically relevant measures, such as sustained attention^5^, memory^13^, craving^14^, PTSD^15^, and social function^16^. While numerous studies have shown that CPM can identify functional connectivity correlates for individual measures (often within specific diagnostic categories), only a handful of studies have examined relationships between functional connectivity and multiple measures in large samples^17–19^. However, these studies used methods different from CPM in their predictive modeling. They also all utilized data from the Adolescent Brain Cognitive Development (ABCD) study. While the ABCD study contains rich fMRI and phenotypic data, brain networks undergo substantial reorganization from childhood to adulthood^20^, potentially impacting the derived brain-behavior models. In addition to ABCD data, one of these studies utilized data from the Human Connectome Project (HCP)^18^ to predict a variety of measures. The HCP also contains high quality fMRI and phenotypic data, but the sample only consists of healthy young adults. Thus, network correlates of clinically-relevant metrics were unable to be identified. The Yale NeuroConnect dataset, described in detail here^21^, provides an opportunity to assess, in a clinically diverse transdiagnostic dataset with extensive fMRI testing (>50 min rest + task), the extent to which a wide range of diagnostically related self-reported clinical symptomatology measures (referred to throughout as ‘clinical measures’ for brevity) and objective cognitive testing instruments have clearly identifiable network correlates and which do not.

The RDoC framework suggests that disordered circuits that affect diagnostically-related subjective symptoms can also impact cognitive abilities. This is corroborated by a large body of evidence indicating that cognitive deficits often co-occur with psychiatric symptoms^22^. However, disordered circuits vary substantially across individuals^23^ with similar symptoms/conditions and may not completely impinge on a specific network related to a particular symptom or cognitive process. Rather, an individual’s disordered circuits are more likely to partially affect multiple networks linked to various symptom and cognitive measures. This phenomenon may underlie the highly variable cognitive deficits seen in psychiatric populations^22,24^. According to this hypothesis, individuals with disordered circuits impinging on networks related to both symptoms and cognition should experience cognitive deficits that co-occur with symptomatology, while individuals with disordered circuits impinging on networks only related to either symptoms should have psychiatric symptoms without cognitive deficits (or vice versa). While similar work has shown that networks associated with mental health measures and cognitive functioning show little overlap^17^, the extent to which connectivity within these overlapping regions contribute to the high amount of variance seen in relationships between subjective measures indexing mental health and objective cognitive assessments remains unknown. To address this question, refinements to computational approaches used for brain-behavior modeling are warranted. Here, we show that CPM can be leveraged to identify circuits that putatively link inconsistencies in cognitive perturbations^22,24^ with high inter-individual variability in network dysfunctions seen in psychiatric populations^23^.

This study has two primary goals. First, using a comprehensive testing battery administered outside the magnet, we determine which of the many clinical and cognitive measures yield CPM models with high prediction strength. This allows us to characterize the extent to which each measure is linked to identifiable brain network correlates while quantifying prediction strength. Furthermore, predicting an array of measures enabled us to address the extent to which measurement noise and other distribution characteristics influence prediction strength, which are important considerations when interpreting brain-based predictive models^25^. Second, we examine the extent to which connectivity in circuits shared between networks associated with both clinical and cognitive measures can predict cognitive performance. With relevance to precision medicine, we show that modeling how disordered circuits related to an individual’s symptomatology impinge upon networks supporting cognition can help explain the variability in cognitive deficits that co-fluctuate with symptomatology. This also demonstrates an advancement in predictive modeling approaches by demonstrating that constraining CPM to networks shared across measures in training data can enhance the precision of functional network localization and identify networks supporting relationships between multiple phenotypes.

## Results

In the present study, we utilized a demographically and clinically diverse transdiagnostic population^21^ (*n*=317; demographic data is presented in **Supplemental Table 1**). The dataset consists of fMRI data from 2 resting-state and 6 task runs, as well as questionnaires collected outside the magnet consisting of 7 real-world self-report clinical assessment scales comprised of a total of 63 distinct measures, and 6 neuropsychological tests yielding 28 distinct cognitive performance measures. The clinical assessment instruments were the Adult Temperament Questionnaire (ATQ)^26^, the Behavior Rating Inventory of Executive Function (BRIEF)^27^, the Brief Symptom Inventory (BSI)^28^, the Interpersonal Reactivity Index (IRI)^29^, the Positive and Negative Affect Schedule (PANAS)^30^, the Pittsburgh Sleep Quality Index (PSQI)^31^, and the Perceived Stress Scale (PSS)^32^. The cognitive testing instruments were the verbal fluency, trail making, color-word interference, and twenty questions subtests from the Delis-Kapan Executive Function Scale (D-KEFS)^33^, the letter-number sequencing, symbol search, and cancellation subtests from the Wechsler Adult Intelligence Scales (WAIS)^34^, the vocabulary and matrix reasoning subtests of the Wechsler Abbreviated Scale of Intelligence 2^nd^ Edition (WASI)^35^, the finger windows and list learning subtests of the Wide Range Assessment of Memory and Learning 2^nd^ Edition (WRAML)^36^, the reading subtest of the Wide Range Achievement Test 5^th^ Edition (WRAT)^37^, and the Boston Naming Test 2^nd^ edition (BNT)^38^. The fMRI data was preprocessed with standard preprocessing methods (described in greater detail in **Methods** and Samardzija et al.^21^). The Shen 268-node atlas^39^ was applied to the preprocessed data, parcellating it into 268 functionally coherent nodes, and functional connectivity matrices were derived for each participant/run. The averages of all 8 functional connectivity matrices were computed for each participant and used as input in building the CPMs. CPM was run for all clinical and cognitive measures in a 10-fold cross validation. Prediction strength was quantified as the Spearman correlation (*ρ*) between the observed measurements and the predicted values. Significance was determined by comparing the prediction strengths from 1000 CPM permutations to a null distribution generated from running CPM on 1000 randomly permuted observations. The proportion of null predictions that exceeded the median prediction strength represented the predictive models’ *p*-values. All *p*-values were FDR corrected for multiple comparisons.

### Prediction strength of clinical measures

To determine whether a measure had sufficient prediction strength to yield reliable brain network correlates, measures must have had *pFDR* < 0.05 and prediction strength > 0.1. In neuroimaging, prediction strengths of 0.1 are generally considered weak, but are accurate enough if also statistically significant to yield meaningful insights about the neurobiological features important to these models^40^. Of 63 clinical measures, 38 (60.32%) met our criteria for being predicted strongly enough to have identifiable whole brain functional connectivity networks (**Fig 1**). The other 25 measures failed to yield sufficient prediction strength to continue with further analyses. When examining predictive model performance across scales and measures, several notable patterns emerged. Of the 7 clinical assessment scales, models built with measures related to sleep quality tended to have the highest prediction strength, with 5 of the top 10 strongest predictions belonging to the PSQI. The mean prediction strength across all PSQI measures was *ρ* = 0.170. Measures pertaining to positive affect from the PANAS also yielded strong predictive models, with 5 out of 6 of these measures providing reliable network correlates (mean *ρ* = 0.151). PANAS measures related to negative affect tended to yield slightly lower prediction strength, with 4 out of 7 providing reliable network correlates (mean *ρ* = 0.120). *ρρρρρρ*Prediction strength values and associated prediction *p*-values can be found in **Supplemental Table 2**, and distributions of clinical measures can be found in **Supplemental Fig 1**.

**Figure 1.**
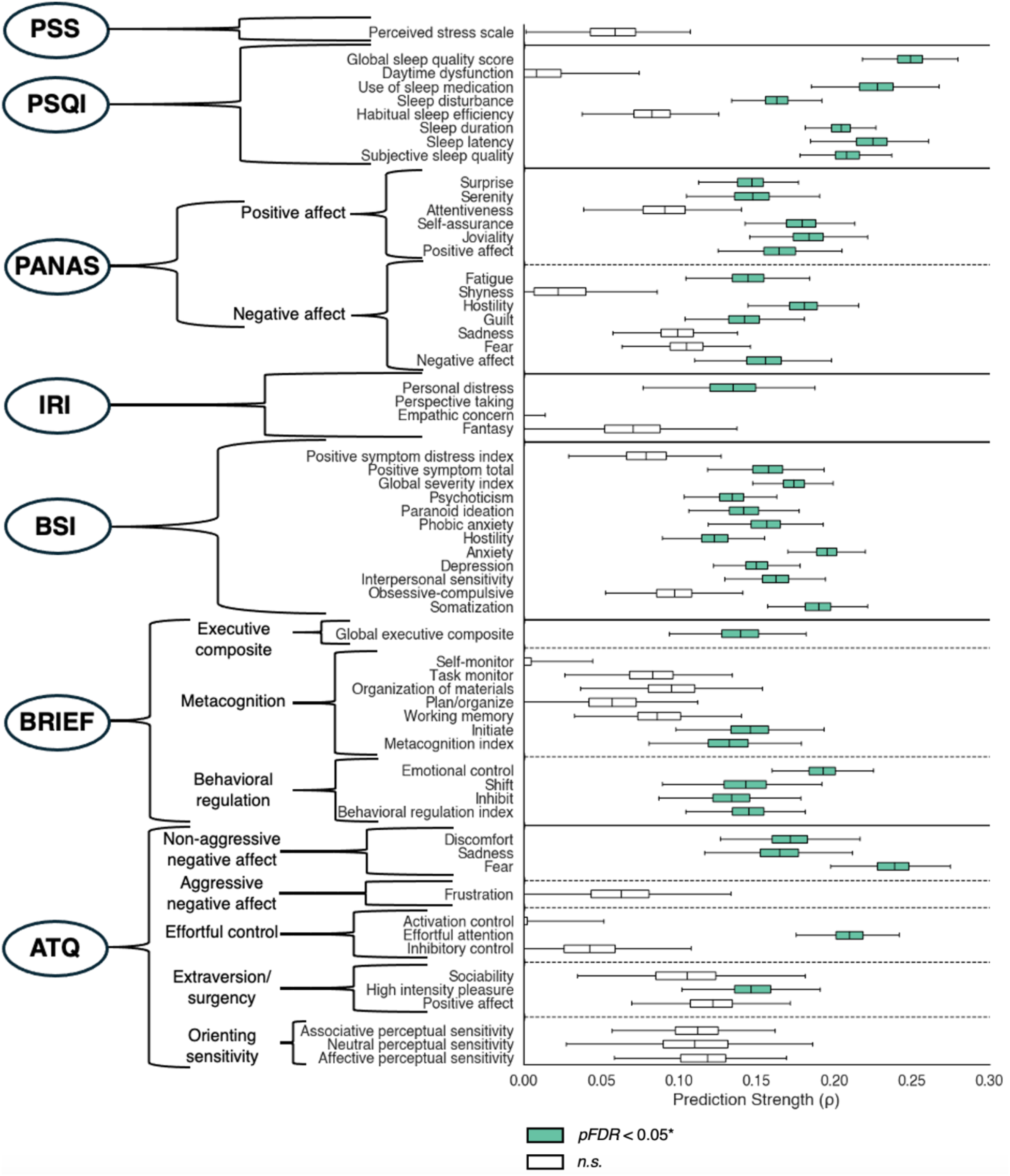
Prediction strength of clinical measures. The boxplots on the right show the prediction strength of each measure and are colored green if significantly predicted by CPM (*pFDR* < 0.05). Measures are grouped according to the instrument they belong to, as indicated by the encircled bolded acronym to the left of the outermost brackets, and the solid horizontal black lines between the boxplots. Measures within instruments were grouped into subdomains if applicable, which are indicated by the middle-level of brackets and the dashed black lines between the boxplots.

Interestingly, there were 4 separate instances where the same measures quantified by different assessment scales yielded widely different predictions. The prediction strength for ‘fear’ was significantly higher when measured by the ATQ than when measured by the PANAS (*ρ* = 0.239 vs *ρ* = 0.105; *p* < 0.001), as was ‘sadness’ (*ρ* = 0.165 vs *ρ* = 0.099; *p* < 0.001). PANAS ‘hostility’ was better predicted than BSI ‘hostility’ (*ρ* = 0.181 vs *ρ* = 0.123; *p* < 0.001). Furthermore, the prediction strength for ‘positive affect’ was higher when measured as an index of all measures related to positive affect by the PANAS than when assessed as a single measure by the ATQ (*ρ* = 0.164 vs *ρ* = 0.122; *p* < 0.001).

Additional models were run covarying for years of education and income in addition to sex and age. In general, the added covariates did not have a strong effect on prediction strength, as the predictions were strongly correlated with the original models (**Supplemental Fig 2**; *r* = 0.946, *p* < 0.001). The networks resolved were also highly similar (**Supplemental Fig 3**; mean Dice = 0.808 +/- 0.058 stdev). In the interest of examining the effects of zero-inflation on the predictions, models were also re-fit with zero-responders removed. Again, the prediction strengths were similar to the original models (**Supplemental Fig 4**; *r* = 0.794, *p* < 0.001), as were the network derivations (**Supplemental Fig 5**; mean Dice = 0.636 +/- 0.189 stdev). However, removing zero-responses did influence the predictability and thus the networks derived for a few measures (specifically, subjective sleep quality, sleep duration, and use of sleep medication had network similarities of Dice < 0.3).

### Prediction strength of cognitive measures

Nearly all cognitive measures yielded excellent prediction strength by CPM. Successful models were built for 26 of 28 (92.86%) measures from the cognitive panel (**Fig 2**). The matrix reasoning test from the WASI provided the model with the highest prediction strength (*ρ* = 0.428). Predictions of measures from the D-KEFS were highly variable, though all produced (range: *ρ* = 0.159 to *ρ* = 0.374). In general, measures from the twenty-questions (mean *ρ* = 0.288) and trail-making tests (mean *ρ* = 0.320) were the most strongly predicted measures from D-KEFS. Performance measures from the WAIS (mean *ρ* = 0.319), the WRAML (mean *ρ* = 0.263), and the WRAT (*ρ* = 0.302) were generally predicted strongly. Measures from the BNT (mean *ρ* = 0.224) and the D-KEFS verbal fluency test (mean *ρ* = 0.177) were predicted less strongly. The two cognitive measures that CPM did not generate reliable models for (WRAML verbal learning intrusion and BNT number correct following a stimulus cue) had the highest percentage of individuals performing at ceiling (see **Supplemental Fig 6**; 53.16% and 77.19%, respectively). These results highlight that predictive models linking external trait measures to brain connectivity patterns via CPM can be built across a wide range of cognitive metrics, with higher prediction strengths representing greater confidence in the derived network correlates. Prediction strength values, associated prediction *p*-values, and brief descriptions of the cognitive abilities theoretically tested by each measure can be found in **Supplemental Table 3**.

**Figure 2.**
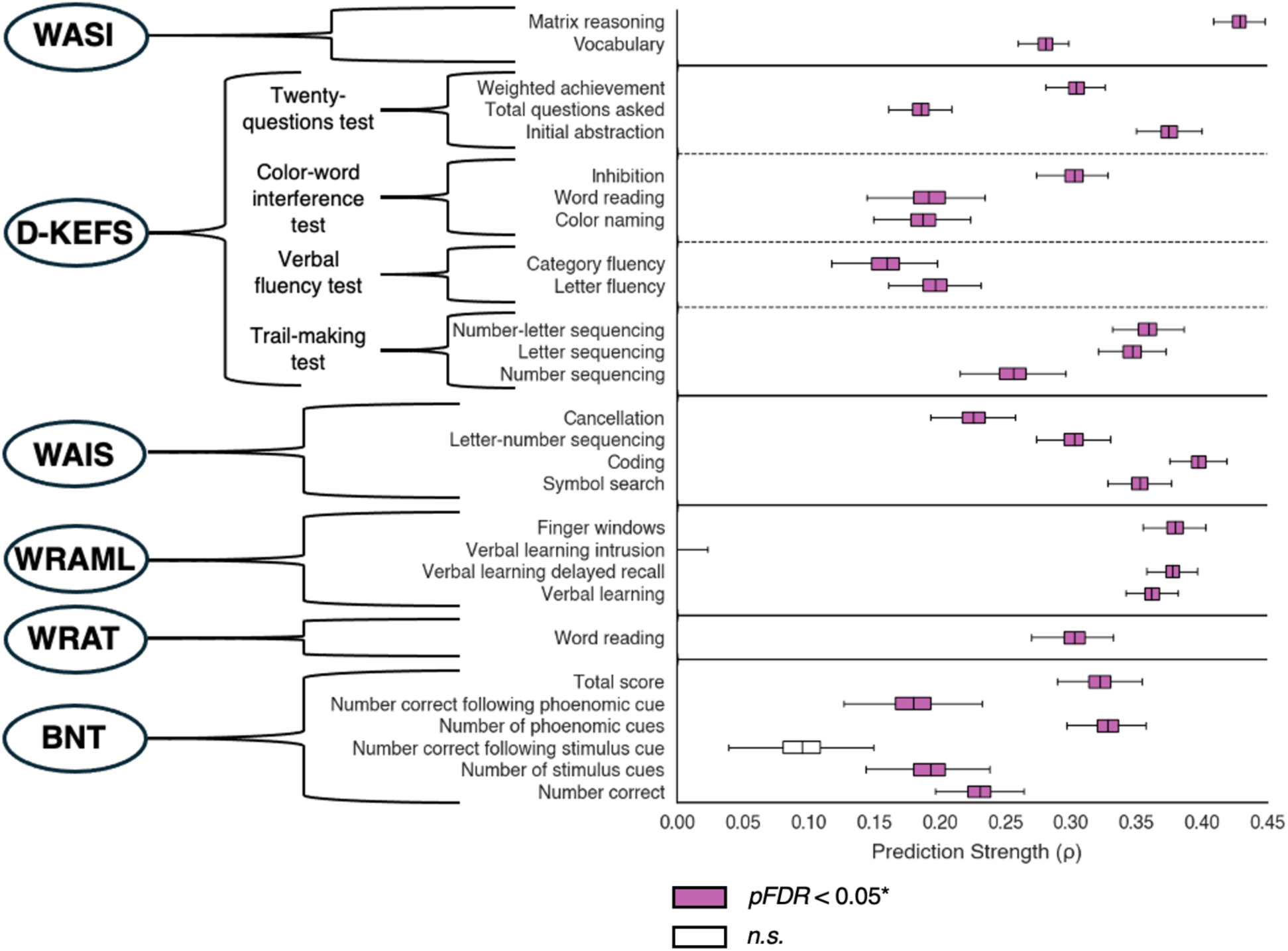
Prediction strength of cognitive measures. The boxplots on the right show the prediction strength of each measure and are colored purple if significantly predicted by CPM (*pFDR* < 0.05). Measures are grouped according to the testing battery they belong to, as indicated by the encircled bolded instrument acronym to the left of the outermost brackets, and the solid horizontal black lines between the boxplots. Measures within the D-KEFS were further subdivided because this battery was composed of subtests that each yielded multiple performance metrics. This is indicated by the middle-level of brackets and the dashed black lines between the boxplots.

Additional models were run covarying for years of education and income in addition to sex and age. The added covariates did not have a strong effect on prediction strength, as the predictions were strongly correlated with the original models (**Supplemental Fig 7**; *r* = 0.967, *p* < 0.001) nor did they substantially change the networks resolved (**Supplemental Fig 8**; mean Dice = 0.719 +/- 0.067 stdev). In the interest of examining the effects of zero-inflation on the predictions, models were also re-fit with zero-responders removed. Similarly, the prediction strengths were highly similar to the original models (**Supplemental Fig 9**; *r* = 0.928, *p* < 0.001), as were the network derivations (**Supplemental Fig 10**; mean Dice = 0.849 +/- 0.105 stdev).

### What drives prediction strength?

In addition to CPM significantly predicting a higher percentage of cognitive than clinical measures, the prediction strength of the cognitive measures that were successfully modeled were higher than the clinical measures’ (**Supplemental Fig 11**; cognitive measures’ mean prediction strength: *ρ* = 0.289; clinical measures’ mean prediction strength: *ρ* = 0.168; *p* < 0.001). This highlights that CPM yields more accurate and robust functional network correlates for cognitive than for clinical measures.

We next sought to understand the factors that influence model performance leading to better prediction strength for some measures compared to others, noting that the functional connectivity data remained constant across all models. Due to the differences in prediction strength between clinical and cognitive assessments, we examined how prediction strength varied across instruments and subdomains tested in clinical and cognitive assessments separately. One-way ANOVAs revealed that prediction strength strongly varied as a function of instrument/subdomain for clinical (**Fig 3a**; *F*(13,62986) = 3059.98, *Cohen’s F* = 0.795, *p* < 0.001) and cognitive measures (**Fig 3b**; *F*(13,27986) = 1194.96, *Cohen’s F* = 0.745, *p* < 0.001).

**Figure 3.**
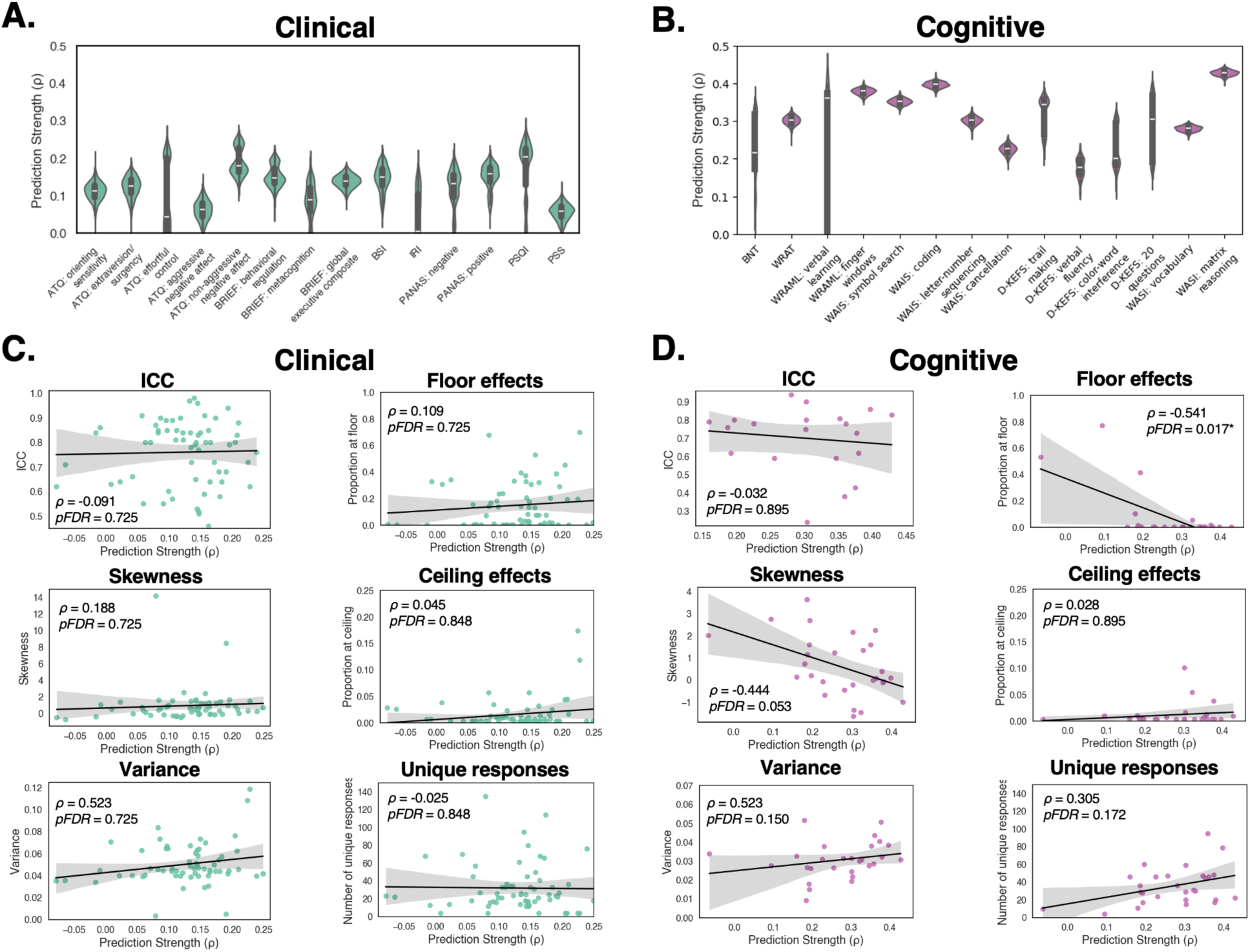
Prediction strength was driven by phenotype measured. **A)** Clinical predictions varied significantly by instrument and subdomain (one way ANOVA: *p* < 0.001). **B)** Cognitive predictions varied significantly by tests and subtests (one way ANOVA: *p* < 0.001). **C)** In clinical measures, prediction strength was unrelated to test-retest reliability (ICC) (top left), skewness (middle left), normalized response variance (bottom left), floor and ceiling effects (top and middle right), and the number of unique responses (bottom right). **D)** In cognitive measures, prediction strength was significantly related to floor effects (top right) and was trending towards being significantly related to skewness (middle left). Prediction strength was unrelated to test-retest reliability (ICC) (top left), the normalized response variance (bottom left), ceiling effects (middle right), and the number of unique responses (bottom right).

Recent work questioned the extent to which brain-based predictive modeling is influenced by measurement noise^25^. To address this, we tested whether the predictive accuracy on these measures was related to their intraclass coefficients (ICCs) reported by the manuals and seminal papers describing each of the testing instruments. We also examined whether model performance was related to other characteristics of the measures’ data (skewness, variance, proportion responding at floor/ceiling, and the number of unique responses). Clinical prediction strength was not correlated with ICC in the measures with reported ICC values (*ρ* = 0.091, *pFDR* = 0.725), nor was it related to the measures’ skewness (*ρ* = 0.188, *pFDR* = 0.725), the variance of the responses (*ρ* = 0.150, *pFDR* = 0.725), the proportion responding at floor (*ρ* = 0.109, *pFDR* = 0.725) and ceiling (*ρ* = 0.045, *pFDR* = 0.848), and the number of unique responses (*ρ* = −0.025, *pFDR* = 0.848) (**Fig 3c**). Similarly, cognitive prediction strength was not correlated with ICC in the measures with reported ICC values (*r* = −0.122, *pFDR* = 0.895), nor was it correlated with response variance (*ρ* = 0.342, *pFDR* = 0.150), ceiling effects (*ρ* = 0.028, *pFDR* = 0.895), or the number of unique responses (*ρ* = 0.305, *pFDR* = 0.172). However, prediction strength of cognitive measures was significantly correlated with floor effects (*ρ* = −0.541, *pFDR* = 0.017*) and was trending towards being significantly correlated with the distribution’s skewness (*ρ* = −0.444, *pFDR* = 0.053) (**Fig 3d**).

### Network similarity across measures

Since disordered circuits could impinge upon both clinical and cognitive networks, we examined the degree of overlap between predictive networks for all clinical and cognitive measures (**Fig 4**). This was achieved by sorting the predictive networks based on similarity using an agglomerative hierarchical clustering algorithm. As expected, the similarities between the measures’ networks mirrored the correlational structure of the measures themselves since the networks are derived from the measures. There was a nearly identical correspondence between these networks and the networks’ Dice similarities when using an unthresholded edge selection approach (*r* = 0.99), indicating that the edge selection threshold that was applied likely did not impact overall network similarity structure.

**Figure 4.**
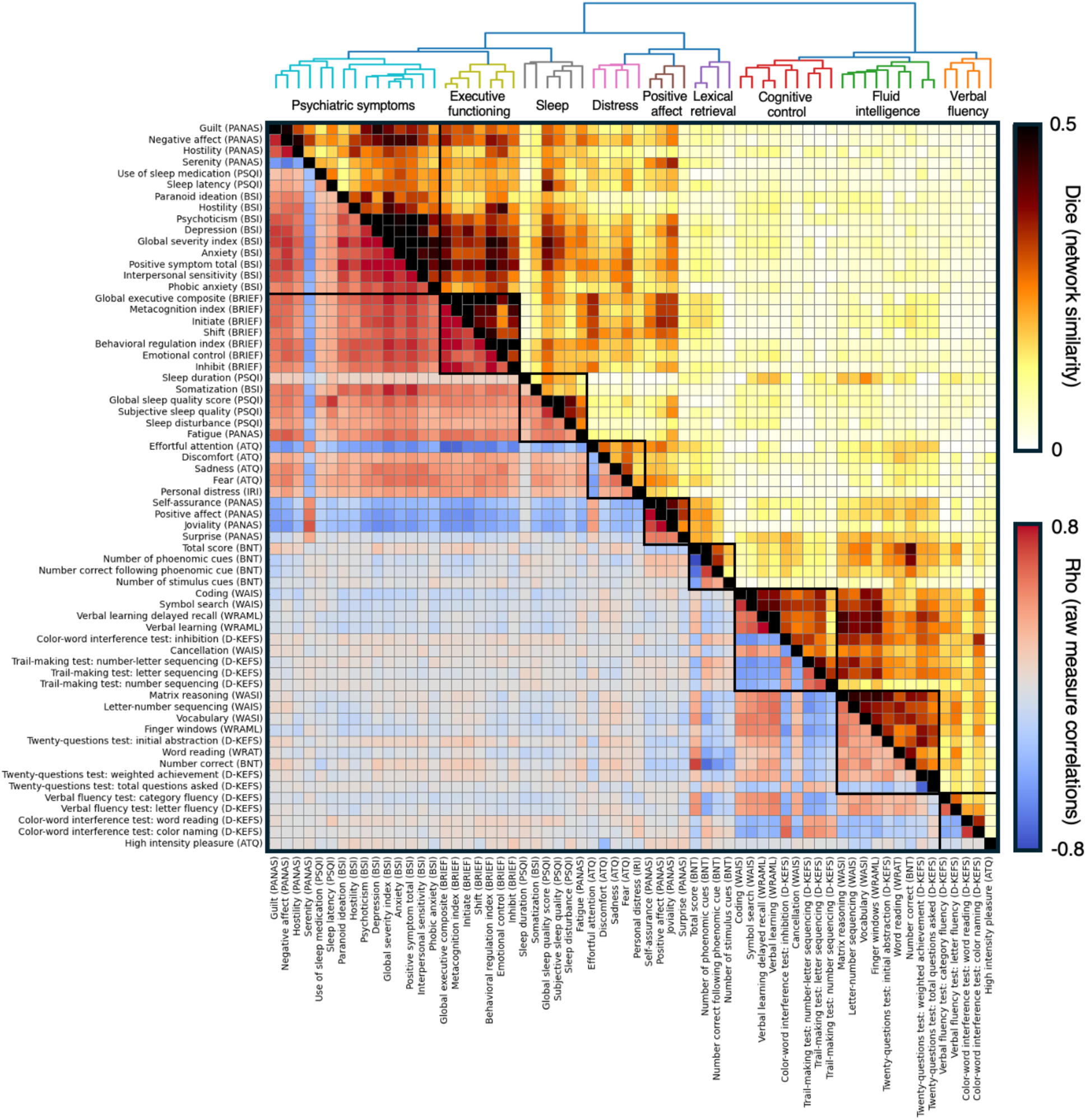
Relationships between measures. The measures’ predictive networks were sorted according to their Dice similarities by an agglomerative hierarchical clustering algorithm (upper triangle). A dendrogram displayed above the matrix highlights how the measures grouped together based on their network similarity. Descriptive labels were given to the 9 clusters and displayed underneath the dendrogram. The measures belonging to each cluster were surrounded black borders on the matrix and colored differently on the dendrogram. The clinical measures were displayed on the top/left and the cognitive measures were displayed on the bottom/right. The network similarity was mirrored by the correlational structure between the raw measures (lower triangle).

In general, the clinical measures were most related to other clinical measures, while the cognitive measures were most related to other cognitive measures. The only exception was high intensity pleasure (ATQ), which was more closely related to cognition than to other clinical measures. The Dice similarity between clinical-cognitive networks was generally much lower than between clinical-clinical (unpaired t-test: *t*(788.66) = −31.68, *p* < 0.001) and cognitive-cognitive networks (unpaired t-test: *t*(389.11) = −23.39, *p* < 0.001). Additionally, after adjusting for multiple comparisons using an FDR correction, only 101 pairs of clinical and cognitive measures were significantly correlated (*pFDR* < 0.05), and their effect sizes were typically weak (maximum *ρ* = 0.234, minimum *ρ* = −0.227). The significantly correlated pairs of clinical and cognitive measures are highlighted in **Supplemental Fig 12**. Together, this suggests that brain networks that vary as a function of cognitive performance are largely distinct from those networks that vary systematically with clinically relevant symptomatology.

We then grouped the brain networks predicting clinical and cognitive measures into 9 clusters based on the hierarchical clustering results. This number was determined using the elbow method^41^ (**Supplemental Fig 13**), which is a commonly used approach for determining an appropriate number of clusters to group data into. These clusters were selected to visualize how the measures grouped together based on network similarity, and to compare the regions important in driving CPM predictions across clusters, rather than to make a claim about how many clusters exist in this dataset. The patterns that the CPM networks loaded onto each of the 9 clusters corresponded to what the measures theoretically assess. Of these 9 clusters, 5 were composed of clinical measures and 4 were composed of cognitive measures. For discussion purposes, each of the clusters were given a name that generally reflects the broad measures composing them. The cluster labels are displayed underneath the dendrogram in **Fig 4**, which align with the sorted measures displayed in the matrix below. It is important to note that these names do not encompass all the behavioral phenotypes present in each cluster. Additionally, since they were not orthogonal, important features were often shared across clusters. For example, cognitive control abilities were likely important to performing all cognitive tests, not just the tests in the ‘cognitive control’ cluster. This phenomenon is reflected by the substantial overlap in the network features between clusters.

### Localizations of networks important to predictions

We examined the contributions of specific networks to prediction strength of clinical measures using the canonical Yeo17 networks as a reference^42^ (**Fig 5a**). For identifying the nodes and edges important to predictions across clusters, we compared positively predicting network features from the positive affect cluster to negatively predicting network features from the 4 other clinical clusters (and vice versa) because these cluster’s measures were anticorrelated. Doing so allowed us to effectively put the brain correlates of measures where higher scores indicated better mental health on the same scale as measures where higher scores indicated worse mental health. The nodes driving the prediction strength for the clinical measures displayed substantial overlap across the clinical measures’ clusters for both positively (mean pairwise Jaccard similarity = 0.252 +/- 0.088 SD, one-tailed *p* < 0.001) and negatively predicting nodes (mean pairwise Jaccard similarity = 0.188 +/- 0.068 SD, one-tailed *p* < 0.001). Examining the top 10% highest degree nodes in each cluster’s predictive networks revealed that the thalamus was the only region important to positively predicting every cluster of clinical measures, and only the cerebellum contained nodes that were important to negatively predicting each cluster of clinical measures. A full list of the top 10% highest degree positive and negative predicting nodes for each clinical measure cluster can be found in **Supplemental Tables 4 and 5**. Many of the same nodes that were important in positive predictions were also implicated in negative predictions (Jaccard similarity = 0.500, one-tailed *p* < 0.001). Furthermore, examining the top 10% of canonical network pairs where the predictive edges were most concentrated showed that limbic A to brainstem connectivity was important to all positive predictions, while within somatomotor A/B and somatomotor A to dorsal attention B were important to negative predictions across all clinical measures’ clusters.

**Figure 5.**
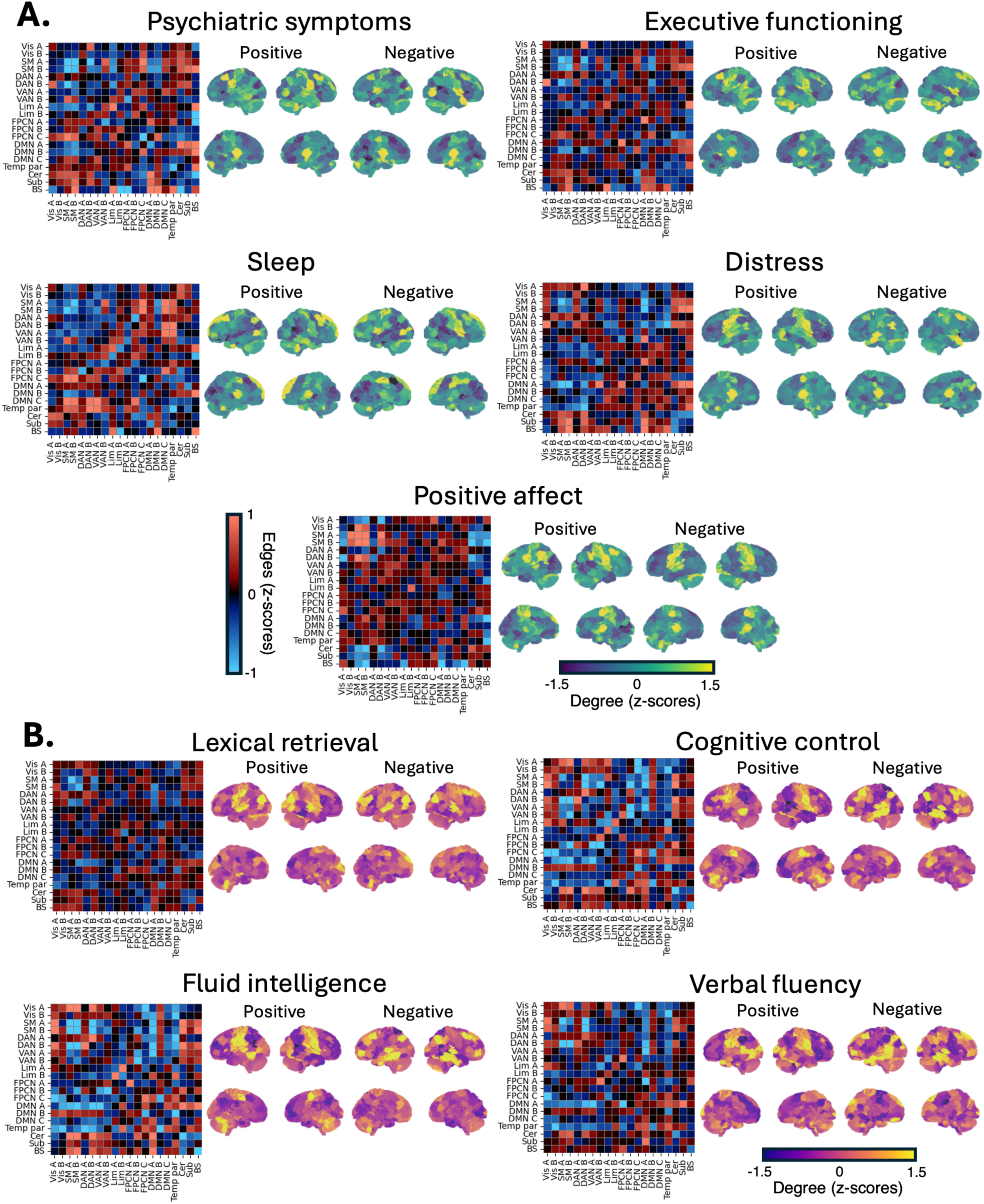
Brain networks driving predictions. **A)** The clusters summarizing networks important to predicting clinical measures are displayed with the descriptive labels ‘psychiatric symptoms’, ‘executive functioning’, ‘sleep’, ‘distress’, and ‘positive affect’. **B)** The clusters summarizing networks important to predicting cognitive measures are displayed with the descriptive labels ‘lexical retrieval’, ‘cognitive control’, ‘fluid intelligence, and ‘verbal fluency’. For each cluster, degree plots highlight the important nodes for predictions across all measures, and matrices highlight the distribution of edges connecting canonical networks. Networks are labeled according to the Yeo 17 network definition, plus cerebellar, subcortical, and brainstem regions.

Similarly, the networks supporting prediction strength in the cognitive measures were assessed (**Fig 5b**). As with the clinical measures, we flipped the sign of the cognitive measures’ predictive networks before comparing across measures to ensure that all positively predicting networks were indicative of better performance on the task, and not just a higher score, since sometimes a higher score was undesirable. The nodes contributing to predicting cognitive measures displayed substantial overlap across the cognitive measures’ clusters for both positively (mean pairwise Jaccard similarity = 0.197 +/- 0.092 SD, one-tailed *p* < 0.001) and negatively predicting nodes (mean pairwise Jaccard similarity = 0.210 +/- 0.118 SD, one-tailed *p* < 0.001). Despite this, examining the top 10% highest degree nodes in each cluster’s predictive networks showed that only nodes within the visual A and somatomotor B networks were positively associated with cognitive performance and only one node in the temporal parietal network was negatively associated with cognitive performance across all clusters. A full list of the top 10% highest degree positive and negative predicting nodes for each cognitive measure cluster can be found in **Supplemental Tables 6 and 7**. There was significant overlap between the nodes important to positively and negatively predicting cognitive measures (Jaccard similarity = 0.294, one-tailed *p* < 0.001), and these nodes often were in the cerebellum. However, this overlap was less pronounced than it was in the clinical measures (Jaccard similarity = 0.294 vs 0.500). The nodes that were only important to positive predictions were typically located toward the bottom of the functional gradient in lower-level unimodal cortex, while nodes that were only important in negative predictions were frequently localized near the top of the functional gradient in higher-level transmodal cortical regions^43^ (see **Supplemental Fig 14**). Examining the top 10% of canonical network pairs where the predictive edges were most concentrated showed that dorsal attention A to dorsal attention B, default mode A to cerebellum, limbic A to limbic B, somatomotor A/B to cerebellum, and visual A to somatomotor B network connectivity was important to positive predictions across all cognitive measures’ clusters. Furthermore, dorsal attention B to default mode A, default mode B to temporal parietal, somatomotor A/B to dorsal attention B, somatomotor B to default mode A, and within somatomotor A network connectivity were important for driving negative predictions across all cognitive measures’ clusters.

### Predicting cognition from impinging circuits within clinical networks

In **Fig 4** we highlighted that the correlations were relatively weak, and network overlaps were sparse between clinical and cognitive measures. This finding suggests that cognitive alterations that co-occur with clinical symptomatology are highly variable between individuals. Given that dysfunctional circuits are heterogeneously distributed in individuals with similar symptoms/conditions^23^, we hypothesized that the precise localization of an individual’s disordered circuitry can be inferred from assessing how their clinical networks impinge upon networks supporting cognitive performance. Therefore, we sought to determine the extent to which circuits that span both systems explain variance in cognitive deficits that co-occur with clinical measures. This was achieved by identifying the edges that were related to both clinical and cognitive measures for all possible clinical-cognitive measure pairings within the training folds of the CPM procedure and then predicting cognitive measures individually from these constrained sets of overlapping edges (**Fig 6a**). Importantly, these models were only trained on edges that predicted poor symptomatology and poor cognition (as opposed to shared edges predictive of poor symptomatology and good cognition; more details in **Methods**).

**Figure 6.**
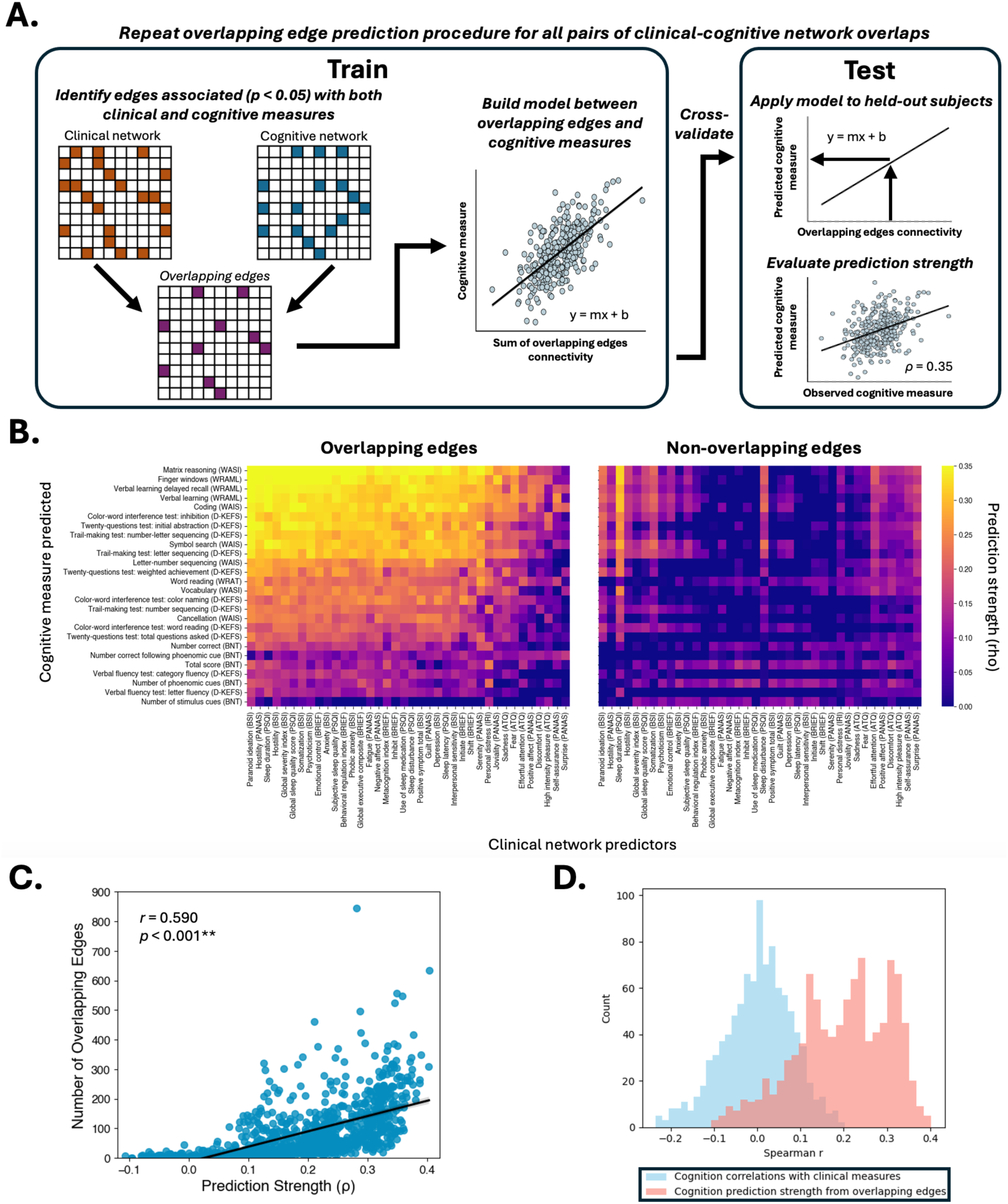
Predicting cognitive deficits from clinical-cognitive network overlaps. A) An overview of how the shared networks were identified and how predictive models were generated. Within each fold of a 10-fold cross validation, edges were identified that were significantly correlated (*p* < 0.05) in the training data with each behavioral measure for all 38 clinical measures and 26 cognitive measures. For every possible pairwise combination of clinical and cognitive measures (38 x 26 = 988 in total), the edges that were related to both measures in the training data were selected for constructing brain-behavior models. These models were then applied to the held-out testing data to predict the cognitive measures. This approach confined the cognitive measures’ predictive networks to just the edges that were impinged upon by the clinical measures’ networks. B) Prediction strength from overlapping edges. The cognitive measures were predicted (y-axis) from the edges in the clinical measures’ networks (x-axis) that their networks overlapped with in the training data (left). Predictions that were not significant (*pFDR* > 0.05) were artificially set to 0 for visualization purposes. To demonstrate the specificity of this effect, cognitive measures were also predicted from edges in the clinical measures’ networks that did not impinge upon cognitive networks in the training data (right). C) There was a significant relationship between the number of overlapping edges and prediction strength *(r* = 0.590, *p* < 0.001**). D) The prediction strength from the overlapping networks tended to be much stronger than the correlations between the clinical and cognitive measures’ values (paired t-test: *p* < 0.001**).

Connectivity in the overlapping edges significantly predicted (*pFDR* < 0.05) cognition 777 times out of 988 total pairs of clinical-cognitive network overlaps (38 clinical x 26 cognitive measures; **Fig 6b**). The cognitive measures were plotted in descending order according to how strongly they were predicted by impinging clinical networks. The 5 cognitive measures that were predicted the strongest were matrix reasoning, finger windows, verbal learning delayed recall, verbal learning, and coding. The 5 impinging clinical networks that predicted these measures most strongly were the paranoid ideation (BSI), hostility (PANAS), sleep duration (PSQI), hostility (BSI), and the global severity index (BSI) networks. As an additional control analysis, models were trained on edges from clinical measures’ networks that did not overlap with cognitive networks and were tested to predict cognitive measures. Importantly, the number of training edges selected for the control models were equal to the number of edges that overlapped between clinical and cognitive networks. Of these 777 significant predictions from overlapping edges, 762 were significantly stronger than cognitive predictions from an equal number of non-overlapping edges within symptom networks (*pFDR* < 0.05). The only instances where the overlapping edge predictions were not significantly stronger than the non-overlapping edge predictions occurred in the some of the weakest overlapping edge predictions (**Supplemental Fig 15**).

The number of overlapping edges was strongly correlated with the prediction strength (**Fig 6c**; *r* = 0.590, *p* < 0.001), indicating that the more widespread the impingements of clinical networks onto cognitive, the more likely that connectivity in these circuits predicted cognition. However, there were 153 instances where cognitive measures were strongly predicted (*ρ* > 0.3) by overlapping networks that contained <100 edges. Furthermore, the prediction strengths yielded by these constrained sets of overlapping edges tended to be much stronger than the correlations between the clinical and cognitive measures’ values (**Fig 6d**; *t*(987) = −59.18, *p* < 0.001), highlighting that shared circuits between clinical and cognitive measures can predict cognition even when the measures themselves are uncorrelated. Taken together, these results suggest that disordered circuits that impinge upon networks supporting cognition account for a significant amount of the variance in cognitive deficits co-occurring with symptomatology.

Finally, we examined where the location of network overlaps important to explaining the variability in clinical-cognitive relationships were. To achieve this, the number of times that each edge was predictive across the 777 overlapping network pairs was identified by summing the binarized predictive network masks. Most predictive edges only overlapped between clinical and cognitive networks a small proportion of times (441 out of 777 overlapping predictive networks shared <50 edges). This was to be expected given the diversity of clinical and cognitive measures assessed. Nevertheless, a subset of edges that were repeatedly predictive across numerous overlapping clinical-cognitive network pairs. Specifically, 291 edges were predictive >50 times, and 96 were predictive >100 times (**Fig 7a**). To examine how these predictive overlapping edges were distributed across the brain, the number of times that overlapping edges were predictive was averaged across all edges within each set of canonical network-network pairs (**Fig 7b**). This way, predictive edges that overlapped more frequently were weighed more heavily.

**Figure 7.**
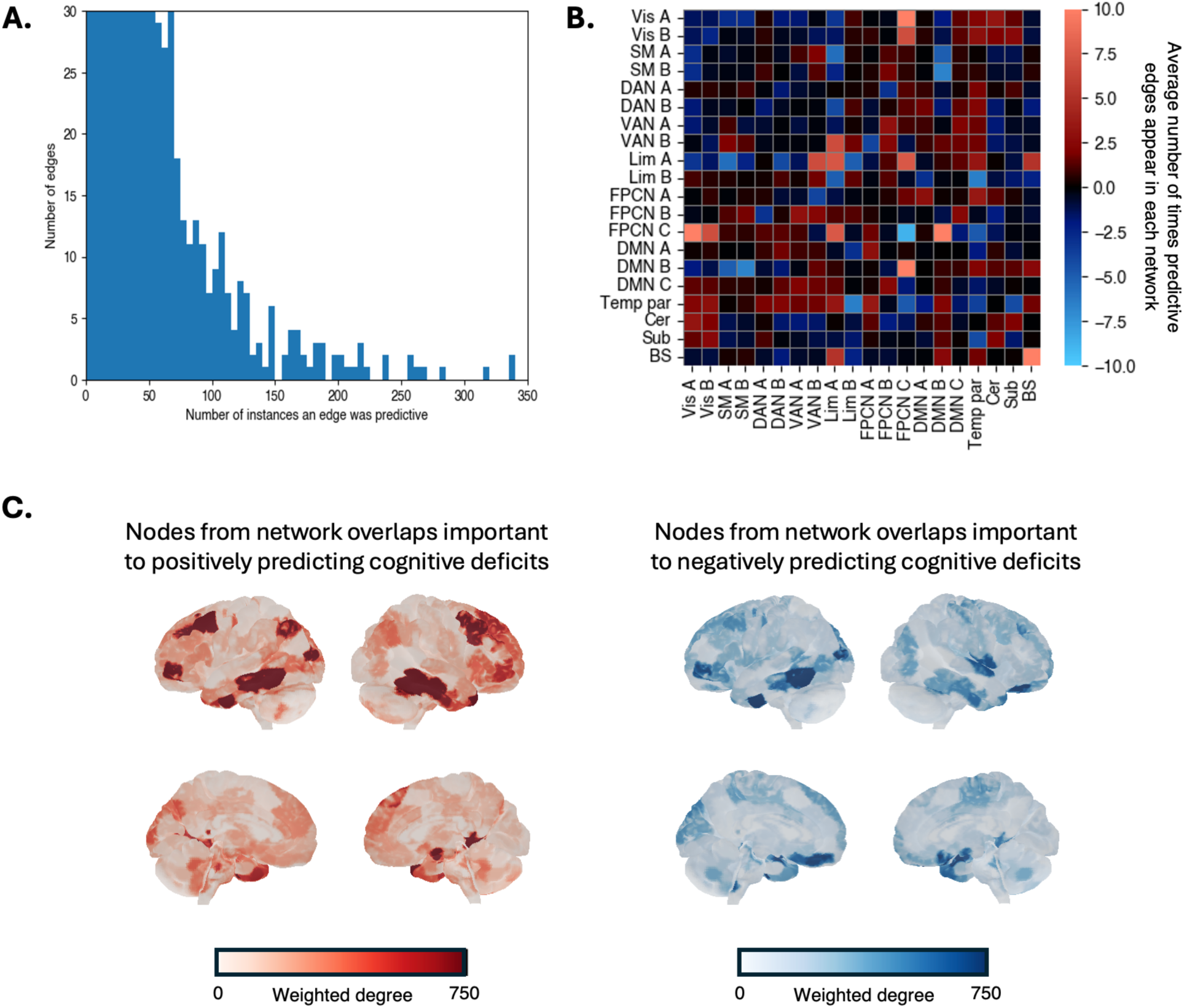
Localizations of overlapping edges that are frequently predictive of cognitive deficits. A) Histogram of how often overlapping edges are predictive of cognitive deficits. Most predictive edges only overlapped between clinical and cognitive networks a small proportion of times (441 out of 777 overlapping predictive networks shared <50 edges). However, there were edges that were frequently implicated in network overlaps. B) A summary of the canonical networks that the overlapping edges were distributed across. The frequency that overlapping edges were predictive was averaged across all edges within each set of canonical network-network pairs. Therefore, edges that often overlapped were given greater importance. Networks were labeled according to the Yeo 17 network definition, plus cerebellar, subcortical, and brainstem regions. The overlapping edges that most frequently positively predicted cognitive deficits were distributed between the frontoparietal control C to default mode B, visual A to frontoparietal control C, within brainstem, limbic A to frontoparietal control C, and within the limbic A network. The overlapping edges that most frequently negatively predicted cognitive deficits were distributed within the frontoparietal control C network. Negatively predicting edges were given negative values for plotting purposes. C) Surface plots displaying the overlapping edges’ nodal degree (weighted by the number of times a predictive edge overlapped). The 5 nodes most important to positively predicting cognitive deficits were in the left dlPFC, right ventral occipitotemporal cortex, bilateral posterior inferior temporal gyrus, and right anterior inferior temporal gyrus. The 5 nodes most important to negatively predicting cognitive deficits were in the left anterior inferior temporal gyrus, bilateral basal ganglia, right orbitofrontal cortex, and left posterior inferior temporal gyrus.

The overlapping edges that most frequently positively predicted cognitive deficits were distributed between the frontoparietal control C to default mode B, visual A to frontoparietal control C, within brainstem, limbic A to frontoparietal control C, and within the limbic A network. The overlapping edges that most frequently negatively predicted cognitive deficits were mostly distributed within the frontoparietal control C network. On average, edges within each of these network-network pairs appeared in overlapping predictive models >7 times. Examining the highest degree nodes within these overlapping networks revealed that the 5 nodes most important in the positive predictive model (edge strength increased with deficit score) for cognitive deficits were in the left dorsolateral prefrontal cortex, right ventral occipitotemporal cortex, bilateral posterior inferior temporal gyrus, and right anterior inferior temporal gyrus. The 5 nodes most important in the negative predictive model (edge strength decreased with increasing deficit score) for cognitive deficits were in the left anterior inferior temporal gyrus, bilateral basal ganglia, right orbitofrontal cortex, and left posterior inferior temporal gyrus. Lists of these nodes’ indices, their centroids’ MNI coordinates, and their canonical network assignments can be found in **Supplemental Table 8**. Overall, these results highlight that performing CPM on sets of edges shared between measures in the training data can yield much more confined solutions for mapping networks related to cognition. Furthermore, this approach revealed that circuits distributed within and between higher order networks are related to cognitive deficits that co-occur with subjective clinical symptomatology measures.

## Discussion

This study had two primary goals. First, we aimed to comprehensively evaluate which clinical and cognitive measures have clear brain correlates as determined by prediction strength in CPM. The findings showed that, in general, there is a tighter coupling between brain networks and external instruments for cognitive tests compared to clinical measures. We also show that this relationship is not uniform across measures, as prediction strength varied substantially across measures in both classes and was not driven by differences in measurement stability. Our second goal was to test a hypothesis upon which the RDoC framework is based: that disordered circuits that affect clinical symptomatology should also impact cognition. Our results indicated that the networks implicated in clinical symptomatology impinge upon networks that support cognitive functioning, and that these overlapping networks explain a significant amount of variance in the relationships between clinically relevant symptoms and cognitive performance metrics. As might be expected, the networks associated with symptoms do not completely align with any specific cognitive network. This work establishes a framework for how to use brain-behavior modeling to locate functional networks that support and facilitate relationships between multiple behavioral domains.

### Clinical predictions and associated networks

The broad testing of clinical measures that we employed in this study allowed us to systematically evaluate the extent to which a variety of clinical assessment instruments can be linked to functional brain networks. We identified reliable functional connectivity correlates of many clinical measures collected in this dataset (38 out of 63 predicted at *ρ* > 0.1 and *pFDR* < 0.05). Varying prediction strength between instruments highlights that some clinical assessment tools are better suited to capture brain network function than others. Among these, the PSQI, the BRIEF (behavioral regulation measures), PANAS, ATQ (non-aggressive negative affect measures) and BSI all tended to be well modeled in the brain. Conversely, measures from the IRI, BRIEF (metacognition measures), ATQ, and the PSS did not consistently yield reliable brain network models. We also observed that some network models of sleep measures were strongly influenced by zero-responses. This suggests that there were a mix of circuits in the original network models that were driven by asymptomatic patients and circuits linked to sleep pathology. Future studies should seek to further understand the relevance of these differing circuits in clinical populations.

Each of these clinical assessment instruments utilized self-reporting. The subjective nature of self-reports introduces several confounds that could interfere with CPM prediction strength, such as response style differences related to introspection abilities^44^, social^45^ and cultural backgrounds^46^, or differences in how respondents and assessment administrators interpret the questions^47^. Measures that are less prone to these influences are thus more likely to have clear network correlates. Understanding how to refine self-report scales to yield better behavioral predictions will improve the extent to which they reflect network functioning, thereby enhancing their utility in identifying neural generators of clinical dysfunction. Better test instruments combined with digital phenotyping with HER and EMA data in its many different forms will help to improve the efficacy of precision medicine^48–50^. Brain models such as CPM can inform the further development of these assessment tools.

The considerable overlap in the networks important to predicting all clinical measures suggests that CPM was capturing a low-dimensional general factor of psychopathology (p-factor) posited by the HiTOP framework and elsewhere^51,52^. Each cluster’s unique deviations from this general p-factor network, therefore, track network dysfunctions specific to the clusters’ phenotypic makeups. Our results strongly implicate the thalamus and cerebellum as key nodes in this p-factor network. These brain structures and their clinical relevance are commonly overlooked, and researchers have argued that this omission is stifling progress in neuroscience^53^. However, there is evidence suggesting the functional connections involving the cerebellum and thalamus contain clinically relevant information^54,55^. Although not involved in predicting clinical measures across all clusters, nodes within the somatomotor network were also frequently involved in predicting clinical measures. Like the cerebellum and deeper brain structures such as the thalamus, the somatomotor network has also been relatively overlooked in psychiatric research. An emerging body of research has linked somatomotor network dysfunction to clinical pathology^56–59^. Our results add to this mounting evidence implicating the thalamus, cerebellum, and SM network in symptomatology.

### Cognitive predictions and associated networks

We also evaluated a comprehensive battery of cognitive tests to identify the extent to which they reliably reflect brain network function. Nearly all cognitive measures yielded models with good prediction strength (26 out of 28 predicted at *ρ* > 0.1 and *pFDR* < 0.05). More than 50% of participants performed at ceiling for the two measures that were not modeled well (verbal learning intrusion (WRAML) and number correct following a stimulus cue (BNT)). These were also the two cognitive measures with the lowest range of scores (see **Supplemental Fig 6**). These results highlight the importance of utilizing cognitive tests with a sufficiently wide distribution of scores to identify reliable functional network associations of cognition. However, considering that there was not a significant correlation between measurement score variance and prediction strength across all cognitive measures (see **Fig 3**), the distribution of the scores does not account for all of the variance in prediction performance.

Since the cognitive tests utilized objective response time- and accuracy-based performance metrics, they were not susceptible to the biases mentioned previously to the same degree as subjective self-report measures. This was likely a major factor contributing to their prediction strength being substantially higher than the clinical measures’ (clinical measures’ mean *ρ* = 0.168; cognitive measures’ mean *ρ* = 0.289). There are several other noteworthy potential factors influencing this difference in prediction strength. It is possible that the networks supporting cognition are less variable than the ones that support psychiatric symptoms. The instruments used to model symptoms could also simply reflect brain function to a lesser extent than cognitive tests. Finally, the cognitive tasks we utilized could have engaged networks that support cognition to a greater extent than networks related to clinical measures. The clinical measures’ predictions may therefore be boosted from tasks that probe states more closely related to what they are assessing. Any of these possibilities could contribute to this difference in prediction strength. Regardless, the higher prediction strength for cognitive tests reported here is accompanied by greater confidence in the derived brain network correlates.

The networks that predicted cognitive measures also showed considerable overlap with each other, suggesting that these common networks track with the g-factor commonly reported in the literature^60^. Given that all the tests probe a spectrum of cognitive processes, this g-factor network likely reflects a mix of the various cognitive components that interact with each other to facilitate test performance. The nodes important for positively predicting cognitive performance often layed at the opposite ends of the principal cortical functional gradient^43^ from negatively predicting nodes, highlighting a departure from the nature of clinical networks, where the same nodes were often important to both the positive and negative predictive models. Interestingly, this pattern did not apply to cerebellar nodes, as areas of the cerebellum that were involved in the positive predictive networks for cognition were also commonly involved in the negative predictive networks. Furthermore, our findings also strongly implicate the SM network in supporting cognitive function, in line with the recent report that systems supporting higher-order cognition are interdigitated within motor cortex^61^.

### Other factors contributing to prediction strength

Recent work has suggested that measurement stability, as reflected by ICC, is a major component underlying predictive modeling performance^25^. We directly tested this by correlating the prediction strength of clinical and cognitive measures with the ICCs reported by the authors of those scales. We observed effects of instrument/subdomain and response variability on predictions, while an effect of ICC was absent across all measures, including those that failed to model well (see **Fig 3**). This does not rule out the possibility that measurement stability is still a factor contributing to brain-based predictive modeling success. Rather, it suggests that if a measurement is sufficiently stable, we can generate a valid brain-behavior model. In most of the measures we used, the ICC values were high (>0.6). Thus, the influence of this on prediction performance should be minimal, and instead the wide range of responses for any given phenotype should dominate.

Additionally, CPM can directly test how the development of the measures can lead to measurable differences in the extent to which they engage clear brain correlates^62^. As demonstrated here, there were 4 instances where different instruments provided measurements with the same label that purportedly measured the same phenotype but yielded widely different predictions (see **Fig 1**). These findings highlight how the framing of the questions probing symptoms or how index scores composed of multiple related measures can improve the confidence of the network correlates derived from predictive modeling.

### Relationships between clinical and cognitive measures

While a large body of work has demonstrated cognitive deficits in psychiatric populations, the findings here indicate that the relationships between symptom and cognitive measures were weak, likely reflecting the highly variable cognitive alterations within clinical populations^22,24,63,64^. When examining the clinical-cognitive relationships more closely, we observed that significant correlations between clinical and cognitive measures were evenly distributed across clusters, suggesting that these measures might be related through more general p- and g-factor correlations. Although we were interested in identifying brain networks associated with both poor symptomatology and impaired cognition, not all clinical measures were associated with poorer cognitive performance. Better performance on cognitive tests that contained a language component (e.g., the BNT) were correlated with negative affect and depression symptoms. One potential interpretation of this finding could be that enhanced verbal abilities feed into more ruminative thoughts and higher levels of depression symptoms^65^. Interestingly, depression symptoms were also negatively associated with performance on WAIS digit symbol coding, a test that assesses cognitive control. This dissociation suggests further interrogation is needed to more thoroughly understand the various cognitive alterations that co-occur with depression symptoms.

### Shared networks explain variance in clinical and cognitive measure relationships

Examining the intersection of the networks for the clinical and cognitive measures revealed that overlap was sparse. These results extend upon similar recent findings that predictive network features are distinct across mental health and cognitive measures^17^ by revealing that connectivity within these shared networks explains a significant amount of the variance in cognitive deficits that co-occur with self-reported clinical symptomatology. This suggests that connectivity in regions where networks overlap could account for much of the heterogeny in cognitive dysfunction seen in psychiatric populations^22,24,63,64^. This also aligns with the RDoC hypothesis that disordered circuits within an individual likely do not impinge directly on a single network, but instead partially affect multiple networks spanning both symptoms and cognitive measures. It also supports the notion that the precise location and distribution of the disordered circuits relate to an individual’s multivariate symptom profile. Thus, identifying overlapping networks can potentially help inform cognitive intervention personalization in clinical populations, which often assume that symptomatology stems from cognitive deficits^22^ and that cognitive remediation can improve symptoms^66^.

To investigate these overlapping networks, we refined CPM such that models were only trained on edges that were related to both cognitive measures and clinical measures. This step improved the specificity of CPM for network mapping. Many of the cognitive measures utilized here reflect a wide range of abilities that are not necessarily a pure reflection of subjective clinical symptomatology. Some measures, such as matrix reasoning, index abilities like general fluid intelligence (a key component of IQ^67^) that are not inherently reflective of pathology. Although clinical symptomatology has been associated with reductions in fluid intelligence, these associations are generally small in magnitude, indicating that substantial variability in fluid intelligence exists independent of clinical symptomatology^68^. Therefore, if general fluid intelligence (or other aspects of cognition) is affected by psychiatric dysfunction, this likely occurs through a partial perturbation of the relevant circuits, rather than impacting entire systems. Our approach enabled us to confine the cognitive measures’ predictive networks to just the edges that were impinged upon by the networks defined for the clinical measures, allowing us to better understand where in the brain circuits supporting cognition are impacted by symptomatology. This may also be useful in other multivariate modeling frameworks, such as examining how brain networks related to anxiety and depression are impacted by poor sleep quality.

Given that patients who experience cognitive impairment may respond differently to various forms of treatment than unimpaired patients^63,64,69^ and that psychiatric patients often consider cognitive impairments among their primary priorities for treatment^70,71^, identifying the loci of these interactions is important and a potential strategy for developing novel diagnostic biomarkers and precision treatment targets. Examining overlapping networks localization revealed that much of the overlap associated with cognitive dysfunction was distributed between task-positive and task-negative networks (e.g., frontoparietal control to default mode, and frontoparietal control to limbic), while network associated with cognitive performance was mostly limited to the task-positive networks (e.g., frontoparietal control). Other research has shown that a breakdown in the anticorrelation structure between task-positive and task-negative networks is a common feature across several dimensions of psychopathology spanning multiple diagnostic categories^72^. Such findings also have been observed in case-control studies of specific diagnoses. Abnormal connectivity withing the frontoparietal control network and breakdowns in the segregation between the default mode and frontoparietal network have been observed in patients with major depressive disorder^73,74^, which is thought to underlie these patient’s deficits in cognitive control^74^. Similar alterations in the connectivity of these networks have also been observed in obsessive-compulsive disorder^75^, bipolar disorder^76^, and schizophrenia^77,78^. The results we present here corroborate these findings and suggest a mechanistic explanation of how disordered circuitry can impact cognitive deficits related to symptomatology transdiagnostically. Additionally, the left dLPFC was among the key hubs in these overlapping networks. Given that recent findings suggest that individuals with treatment resistant depression who also have cognitive impairments respond better to TMS applied to the left dLPFC than patients with intact cognition^64^, this indicates that this region may be a promising target in precision medicine settings. Future work should therefore further examine the utility of these overlapping circuits in biotyping and guiding personalized interventions.

### Limitations

There are some noteworthy limitations to the current study. First, without an external sample to validate our results, we cannot be certain that the network correlates of various behavioral measures will generalize to unseen data. This is one of the first studies to utilize a large transdiagnostic sample, with both extensive fMRI data (>50 min) and comprehensive phenotyping conducted outside the magnet^21^. Similar data to evaluate whether these results generalize is lacking. Second, some of the studies that we obtained ICC values from were inconsistent in the time intervals between test and re-test, sample sizes, and age groups utilized. These differences could have biased ICC values, potentially obscuring any effects on prediction accuracy. Third, we acknowledge that there are many different methods to determine “optimal” clustering solutions that each typically yield widely different results, and that there is often no clear determination of how many clusters exist^79^. The clustering performed here was solely intended for reducing the dimensionality of the data and showing how similar the measures were to each other based on their network correlates. Their labels were therefore only used to help describe the types of measures composing each cluster. Fourth, the biases introduced via self-reporting reduce the prediction strength of clinical measures and add noise to our network estimates for those measures. However, most instruments available for assessing psychiatric symptoms in clinical contexts rely on self-reporting, and objective assessments are lacking. Fifth, networks associated with clinical measures may be more heterogeneously affected than networks associated with cognitive functioning, especially in a transdiagnostic population that includes many healthy individuals. The asymmetry may underlie the differences in prediction strength between clinical and cognitive measures and should be further explored in populations with less expected heterogeneity in networks associated with clinical measures. Sixth, we have loosely used the terms cognitive and clinical or symptom measures. Some of these objective cognitive tests were designed to ascertain clinically-relevant information regarding cognitive dysfunction, and some of these clinical measures were designed to assess subjective, real-world cognitive functioning. None of these measures are purely cognitive nor purely clinical and the categorical distinction here is not binary. As the data shows, these are not independent systems. Nevertheless, it helps to frame the discussion in this context which is commonly used in the literature. Finally, our results beg a deeper investigation into the relationship between symptoms and cognition. We have assumed here that we are modeling fixed traits, but it is well known that brain-state can strongly influence findings in network neuroscience^80^. Symptoms and cognition wax and wane in both healthy and clinical populations. Understanding how these models may vary, and how the relationship between symptoms and cognitive variables interact over time could be very informative in clinical contexts^80^.

## Methods

### Participants

The sample reported in this study includes data from 317 demographically diverse participants with at least 50 minutes of high-quality functional MRI data and extensive phenotyping outside of the magnet. Participants were recruited from advertisements broadly distributed throughout the New Haven community and referrals from Yale clinics. This transdiagnostic population experienced a wide range of symptoms and symptom severities, often with multiple psychiatric diagnoses. All participants provided written informed consent in accordance with a protocol approved by the Yale IRB. A more thorough description of this dataset can be found elsewhere^21^.

### Imaging acquisition protocol

The imaging data was collected at Yale on a 3T Siemens Prisma scanner with a 64-channel head coil. A high-resolution T1-weighted MPRAGE (TR= 2,400 ms, TE = 1.22 ms, flip angle = 8°, voxel size 1 mm^3^) was acquired for each participant. The functional data were obtained using a multiband EPI sequence (TR = 1,000 ms, TE = 30 ms, flip angle = 55°, slice thickness = 2 mm, multiband factor = 5). A total of 6-task and 2-resting-state runs were acquired, each lasting 6 minutes and 49 seconds. The first and last functional scans were resting-state runs. The six task runs were all designed as event-related designs and comprised a working memory (n-back), an inhibition (stop signal task), a decision making (card guessing), an emotional perception (eyes), a continuous performance task (the gradCPT), and a more naturalistic movie watching task. The task order was randomized and counterbalanced across participants. There was no block structure to the tasks and they were designed as state-manipulation tasks for functional connectivity analyses unlike a typical task paradigm designed for conventional GLM analyses. We ensured that participants understood task instructions and received practice. Responses were recorded on a button box. Analyses were restricted to participants who completed all fMRI scans runs.

### Functional MRI data processing

Skull stripping of the structural scans was done using an optimized version of the FMRIB’s Software Library (FSL) pipeline^81,82^. Motion correction was done using SPM12^83^. Nonlinear registration of the MPRAGE to the MNI template was performed using BioImage Suite^84^, and linear registration of the functional to the structural images was done through a combination of FSL^85^ and BioImage Suite. The remaining preprocessing steps were performed in BioImage Suite, including global signal regression, high-pass and low-pass filtering, and regression of motion parameters. All registered data were visually examined to ensure whole-brain coverage, adequate registration, and the absence of artifacts or other quality issues. Subjects were included in the study if they completed all eight fMRI scan runs and had a grand mean frame-to-frame displacement of less than 0.15 mm and a maximum mean frame-to-frame displacement of less than 0.2 mm. Two participants were scanned with a slightly shorter scanning time (25 seconds shorter) and were included in the dataset.

The Shen 268 node atlas^39^ was applied to the preprocessed data, parcellating it into 268 functionally coherent nodes. Pearson correlations of the time series between all node pairs were computed and subsequently z-transformed to generate 8 functional connectivity matrices for each participant. The averages of all functional connectivity matrices were computed for each run/participant. Since individuals’ functional connectomes are more similar to themselves, regardless of the task, than they are to other individuals^3^, the average connectome will still likely contain relevant information about an individual’s trait characteristics. By including connectivity data collected during multiple different cognitive tasks in the averaged connectomes, patterns of task-evoked brain activity that may differentially enhance behavioral prediction were accounted for^6^. Analyses and visualizations were conducted using both MATLAB and python.

### Clinical assessments

A battery of commonly used self-report questionnaires probing a wide range of clinically relevant measures were collected immediately following the MRI scans (same day). They were selected to index aspects numerous constructs from each of the six major domains of basic human neurobehavioral functioning specified by the RDoC framework^1,2^ (negative valence systems, positive valence systems, cognitive systems, social processes, arousal and regulatory systems, and sensorimotor systems). The assessments consisted of the Adult Temperament Questionnaire (ATQ)^26^, the Behavior Rating Inventory of Executive Function (BRIEF)^27^, the Brief Symptom Inventory (BSI)^28^, the Interpersonal Reactivity Index (IRI)^29^, the Positive and Negative Affect Schedule (PANAS)^30^, the Pittsburgh Sleep Quality Index (PSQI)^31^, and the Perceived Stress Scale (PSS)^32^. Two of the questionnaires (BSI and BRIEF) provided both raw and age and sex normalized scores. In this study, we used the raw scores for neuro-behavioral modeling to maintain consistency across clinical measures. Some participants did not complete all clinical assessment scales. They were included in all analyses for which they had data. A detailed breakdown of the number of subjects (*n*s) for each measure can be found in **Supplemental Table 9**.

### Cognitive assessments

A series of neuropsychological evaluations was administered by a trained evaluator to objectively test a wide array of cognitive functions related to the cognitive constructs specified by the RDoC framework^1,2^ (attention, perception, cognitive control, language, declarative memory, and working memory). These were conducted in the same post-imaging session as the clinical assessments. In total, we administered the verbal fluency, trail making, color-word interference, and twenty questions subtests from the Delis-Kapan Executive Function Scale (D-KEFS)^33^, the letter-number sequencing, symbol search, and cancellation subtests from the Wechsler Adult Intelligence Scales (WAIS)^34^, the vocabulary and matrix reasoning subtests of the Wechsler Abbreviated Scale of Intelligence 2^nd^ Edition (WASI)^35^, the finger windows and list learning subtests of the Wide Range Assessment of Memory and Learning 2^nd^ Edition (WRAML)^36^, the reading subtest of the Wide Range Achievement Test 5^th^ Edition (WRAT)^37^, and the Boston Naming Test 2^nd^ edition (BNT)^38^. A few of the cognitive tests provided both raw and age and sex normalized scores. In this study, we used the raw scores for predictive modeling to maintain consistency across all measures. Some participants did not complete all cognitive tests. They were included in all analyses for which they had data. A detailed breakdown of the number of subjects (*n*s) for each measure can be found in **Supplemental Table 10**.

### Predictive modeling

We used CPM^3,10^ to identify functional brain networks that were related to each clinical and cognitive measure separately. Each predictive model was trained and tested in a 10-fold cross-validation, which was repeated 1000 times to obtain a distribution of predictions. Within each training fold, edges were selected if they were significantly correlated (*p* < 0.05) with behavior after controlling for age and sex using partial correlations. Given that cognitive abilities are often correlated with years of education^86^, and many clinical symptoms are often influenced by income^87^, these demographic variables were not controlled for in the main analyses (though they were in a supplemental analysis). Prediction strength was determined by comparing the observed clinical/cognitive measures to the predicted measures using Spearman correlations. Permutation tests were utilized for significance testing. Specifically, the number of null prediction values that were greater than the median prediction for each measure was divided by the number of permutations to derive the *p*-values. The *p*-values were FDR-corrected for 91 comparisons (63 clinical + 28 cognitive measures). Prediction strengths between the same measures assessed by different instruments were formally compared by comparing the difference between the two median prediction strengths to the difference of median prediction strengths from the two distributions predictions after shuffling, bootstrapped 1000 times. The number of instances when the median difference from the shuffled predictions exceeded the true median prediction strength differences was divided by 1000 (the number of bootstraps) to derive *p*-values. Networks for each measure were derived by identifying the edges that were predictive in >50% of permutations. The resulting networks were binarized, with a ‘1’ or ‘0’ indicating an edge’s network membership. In the interest of addressing the extent to which this 50% threshold influenced the overall network mapping structure, networks that were weighted based on the number of times an edge was predictive across permutations were also computed.

### Assessing the effects of instruments and subdomains on prediction strength

We utilized one-way ANOVAs to assess the effect of measurement instruments and subdomains on prediction strength for clinical and cognitive measures separately (**Fig 3a/b**). To do so, prediction strength values from measures within the same clinical assessment subdomain or cognitive testing battery were grouped together and compared to other subdomains/tests. The ATQ, BRIEF, and PANAS inventories explicitly specified that the individual measures belonged to separate subdomains (e.g., PANAS measures were split into ‘positive’ and ‘negative’ affect domains). These separate subdomains were treated as separate categories for this analysis. The BSI, IRI, PSQI, and PSS inventories did not specify how measures could be grouped into separate subdomains. Therefore, all measures within these instruments were categorized as belonging to the same subdomain for this analysis. Several of the cognitive testing batteries were composed of sets of multiple separate cognitive tests. The D-KEFS consisted of the twenty-questions test, the color-word interference test, the verbal fluency test, and the trail making test. Additionally, the WASI was composed of a vocabulary test and a matrix reasoning test, the WRAML was composed of a verbal learning test and a finger windows test, and the WAIS consisted of cancellation, letter-number sequencing, digit-symbol coding, and symbol search tests. These were all categorized as belonging to different subdomains for this analysis, since they were all different cognitive tests. Conversely, all the metrics from the BNT captured various aspects of BNT performance and were thus considered as belonging to the same subdomain.

### Obtaining measurement reliability data

Test-retest reliability (ICC) measures were obtained from the original papers and/or technical and user manuals published for each clinical and cognitive inventory. The BRIEF provided separate ICC data from a normative sample and a mixed clinical/healthy adult population. We used the ICC data from the mixed clinical/healthy adult sample, since it more closely matched our study’s participant population. The original paper validating the ATQ provided ICC data collected from 2 separate studies, one conducted in undergraduate students and the other in adults between 26 and 91 years old (median age = 57 years). We utilized the data from the study conducted in undergraduate students because it provided ICC data for all the ATQ measures that we used here, while the other study did not. Additionally, our sample’s median age (31.17 years) is closer to the median age of typical undergraduate populations. The IRI reported ICC data from males and females separately. We averaged those results for use in the analysis here. There were no ICC data reported for the ‘total questions asked’ measure from the twenty-questions subtest in the D-KEFS, the verbal learning intrusion measure in the WRAML, or for the performance measures on the BNT (except for ‘total score’). Therefore, these measures were omitted from analyses examining the relationship between ICC and prediction performance.

### Calculating distribution characteristics

For each clinical and cognitive measure, skewness, variance, proportion performing at floor and ceiling, and the number of unique responses were calculated and compared to prediction strength. Skewness was calculated using python’s scipy library. Variance was calculated by first performing min-max scaling to normalize and rescale each variable to a range from 0 to 1. The proportion performing at floor and ceiling were calculated by identifying the number of people with the min/max scores and dividing that number by the number of respondents.

### Identifying relationships between measures and calculating network similarity

Spearman correlations were calculated for each pair of measures that CPM significantly predicted, and the derived *p*-values were FDR corrected for multiple comparisons. Network similarity between measures was assessed using Dice similarity coefficients. If there was a positive relationship between two measures, then the Dice coefficient was calculated between the positively predicting network for measure A and the positively predicting network for measure B, and then calculated once more between the negatively predicting network for measure A and the negatively predicting network for measure B. If there was a negative relationship between two measures, then the Dice coefficient was calculated between the positively predicting network for measure A and the negatively predicting network for measure B, and then calculated once more between the negatively predicting network for measure A and the positively predicting network for measure B. A weighted average of these two Dice coefficients was taken and used to examine overall network similarity between measures. This same approach was performed when comparing the unthresholded networks, except weighted Dice coefficients were utilized.

### Comparing network similarity across measures

We sorted the measures based on network similarity in the interest of identifying meaningful commonalities and differences between predictive networks across measures. To achieve this, we input the inverse of this full Dice similarity matrix (i.e., Dice distances) into an agglomerative hierarchical clustering algorithm using Ward’s method. This yielded a dendrogram that sorted the measures based on their predictive networks’ similarities. The measures were grouped into clusters based on the hierarchical clustering results. The number of clusters chosen was determined using the elbow method^41^, which evaluates where the incremental decrease in within-cluster sum of squares (WCSS) begins to slow across increasing values of *k*. The point of maximum curvature (i.e., the “elbow”) was objectively identified using the KneeLocator algorithm^88^. This is a commonly used approach for determining an appropriate number of clusters in a dataset. However, we performed clustering with the intention of reducing dimensionality, showing how the measures grouped based on network similarity, and to compare important features driving CPM predictions across measures, rather than to make any claims about how many clusters exist in this dataset.

To identify the extent to which the same nodes were important to positively and negatively predicting clinical and cognitive measures, we calculated the Jaccard similarity between vectors representing the top 10% highest degree nodes in the positive and negative predictive networks for each cluster. Overlap was summarized as the mean of the pairwise Jaccard values, and one-sided permutation tests (10,000 iterations) were utilized for significance testing. The *p*-values were calculated as the proportion of null replicates that exceeded the observed mean Jaccard similarity.

### Identifying localizations of predictive networks

The binarized positively and negatively predicting networks for measures within a cluster were summed separately and Z-scored. This yielded aggregate positively and negatively predicting networks for each cluster, with edges most important to predictions across measures weighted most heavily. Before summing the networks, we ensured that the directionality of all the measures within a cluster was similar so that networks positively related to symptoms or cognition would not be mistakenly combined with networks negatively related to symptoms or cognitions. For example, if there was one measure in a cluster where a higher score was desirable while lower scores were desirable for the remaining measures, then the positive network for the first measure would be summed with the negative networks for the other measures (and vice versa). The distribution of edges within and between canonical networks was evaluated by summing the edges within each canonical network pair from the aggregated cluster networks. We utilized the Yeo17 networks^42^ as network definitions, plus the cerebellum, brainstem, and subcortical regions. The nodes important to predictions were examined by calculating degree centrality for each node. The degree values were then projected onto the FreeSurfer template brain’s (‘fsaverage’) pial surface, along with cerebellar and brainstem surface reconstructions rendered in Bioimage Suite. These degree plots were created using Python’s nilearn package^89^.

### Predicting cognition from network overlaps

Prior to using clinical and cognitive network overlaps to predict cognitive performance, measures in which a higher score was more desirable were inverted in their direction so that for all measures, a higher score was less desirable. This was done because we were interested in how poor clinical symptomatology related to cognitive deficits. Inverting the scores facilitated the ease by which shared edges that were positively/negatively associated with both clinical and cognitive measures in the training data could be identified. CPM models were subsequently run on all measures within the same training/testing cross validation folds. Within each training fold, for all possible pairwise combinations of clinical and cognitive measures, edges that were positively correlated (*p* < 0.05) with both clinical and cognitive measures (after controlling for age and sex using partial correlations) were summed together with the negative sum of the edges that were negatively associated with both clinical and cognitive measures (also after controlling for age and sex using partial correlations). A linear model describing the relationship between the cognitive measure and the summed edges associated with both the cognitive and clinical measures were constructed. Then, that model was applied to the held-out testing data to predict the cognitive measure. This was performed for all 38 clinical measures and 26 cognitive measures that were previously determined to be modelable by CPM, yielding a total of 988 different cognitive measure predictions from sets of overlapping edges. These 10-fold cross-validation models were repeated 1000 times to obtain a distribution of predictions. Prediction strength was determined by comparing the observed cognitive measures to the predicted measures using Spearman correlations. Permutation tests were utilized for significance testing. Specifically, the number of null prediction values that were greater than the median prediction for each measure was divided by the number of permutations to derive the *p*-values. The *p*-values were FDR-corrected for 988 comparisons. Predictive overlapping networks were derived by identifying the edges that were predictive in >50% of permutations. The resulting networks were binarized, with a ‘1’ or ‘0’ indicating an edge’s network membership.

To demonstrate that the specificity of the overlapping edges to predicting cognitive measures, an additional control analysis was performed in which an equal number of edges to the number of overlapping edges were randomly selected from the clinical networks in regions that did not overlap with the cognitive networks. This was similarly performed in the 10-fold cross validation procedure described above. Prediction strengths from these control analyses were compared to the prediction strengths from the overlapping network predictions by comparing the difference between median prediction strengths of the two sets of predictions to the difference of median prediction strengths from the two sets of predictions after shuffling, bootstrapped 1000 times. The number of instances when the median difference from the shuffled predictions exceeded the true median prediction strength differences was divided by 1000 (the number of bootstraps) to derive *p*-values. The *p*-values were FDR-corrected for 988 comparisons.

### Localizing overlapping edges that predicted cognition

Localization of overlapping edges that were predictive of cognition was achieved by summing the binarized overlapping network masks. By doing this, the frequency that predictive edges overlapped were indexed and weighed more heavily in the localization analyses. These overlapping edge frequency indices were averaged for all edges within each set of canonical network pairs to summarize the how the edges were distributed across networks. Again, the Yeo17 networks^42^ were utilized as network definitions, plus the cerebellum, brainstem, and subcortical regions. In addition to edge distributions across networks, the localizations of the nodes commonly implicated in overlapping edge predictions were examined. To do so, the matrices indexing the frequencies of positively and negatively predicting overlapping edges were summed separately. This yielded two 268-element vectors corresponding to the nodal degree for each node in the Shen268 atlas. The degree centralities of the summed overlapping positively and negatively predicting networks were visualized on the brain surface the same way as before using Python’s nilearn package^89^.

## Acknowledgements

This work was supported by funding from the National Institutes of Health (MH121095 to D.S. and R.T.C. and MH138347 and EB034720 to R.T.C.). AJS is also supported by the National Science Foundation’s Graduate Research Fellowship Program. We would like to thank Xenophon Papademetris for help with data visualization and Sydney Smith for helpful conversations about data interpretation.

## Code availability

The code used in all of the main analyses conducted in this study can be found at: https://github.com/YaleMRRC/YaleNeuroConnect_FullPredictions.git. All dependency files required to run this code are also provided in this GitHub repository (e.g., cortical surface files), as is the relevant input data.

