## Supplemental materials for "Transdiagnostic connectome-based predictive modeling reveals where circuits related to self-reported clinical symptoms impinge upon brain networks supporting cognition"

| | |
|---|---|
| **Sex** | M = 143<br>F = 174 |
| **Age** | mean = 31.17 +/- 11.17 years |
| **Education** | mean = 15.85 +/- 3.12 years |
| **Race** | White (not Hispanic) = 149<br>Black (not Hispanic) = 50<br>Biracial (not Hispanic) = 7<br>White (Hispanic) = 34<br>Black (Hispanic) = 9<br>Biracial (Hispanic) = 8<br>Asian or Pacific Islander = 48<br>Native = 1<br>Other = 11 |
| **English as first language** | Yes = 272<br>No = 45 |
| **Employment** | Full time = 136<br>Part time (regular hours) = 37<br>Part time (irregular hours) = 31<br>Student = 92<br>Retired/disability = 4<br>Unemployed = 14<br>Residential setting = 1<br>Other = 2 |
| **Income** | $0 - $19,999 = 68<br>$20,000 - $34,999 = 60<br>$35,000 - $49,999 = 53<br>$50,000 - $99,999 = 81<br>$100,000 - $249,999 = 43<br>Over $250,000 = 10 |

**Supplemental Table 1. Demographic information.** For each categorical variable, number of participants were reported. Mean values were reported for continuous variables with +/- indicating standard deviations. There were 3 individuals who did not report their household income levels.

| Assessment | Measure | Prediction strength ($\rho$) | P-value (FDR corrected) |
|---|---|---|---|
| ATQ | Activation control | -0.016 +/- 0.028 | 0.548 |
| ATQ | Effortful attention | 0.21 +/- 0.013 | < 0.001 |
| ATQ | Inhibitory control | 0.042 +/- 0.026 | 0.333 |
| ATQ | Affective perceptual sensitivity | 0.118 +/- 0.022 | 0.066 |
| ATQ | Neutral perceptual sensitivity | 0.11 +/- 0.030 | 0.099 |
| ATQ | Associative perceptual sensitivity | 0.112 +/- 0.020 | 0.068 |
| ATQ | Frustration | 0.062 +/- 0.028 | 0.221 |
| ATQ | Fear | 0.239 +/- 0.015 | < 0.001 |
| ATQ | Sadness | 0.165 +/- 0.018 | 0.021 |
| ATQ | Discomfort | 0.171 +/- 0.017 | 0.009 |
| ATQ | High intensity pleasure | 0.146 +/- 0.017 | 0.034 |
| ATQ | Positive affect | 0.122 +/- 0.021 | 0.063 |
| ATQ | Sociability | 0.105 +/- 0.028 | 0.091 |
| BRIEF | Behavioral regulation index | 0.145 +/- 0.015 | 0.020 |
| BRIEF | Inhibit | 0.134 +/- 0.018 | 0.033 |
| BRIEF | Shift | 0.143 +/- 0.021 | 0.031 |
| BRIEF | Emotional control | 0.193 +/- 0.012 | 0.003 |
| BRIEF | Metacognition index | 0.132 +/- 0.019 | 0.047 |
| BRIEF | Initiate | 0.146 +/- 0.019 | 0.034 |
| BRIEF | Working memory | 0.086 +/- 0.021 | 0.123 |
| BRIEF | Plan/organize | 0.057 +/- 0.022 | 0.238 |
| BRIEF | Organization of materials | 0.095 +/- 0.022 | 0.102 |
| BRIEF | Task monitor | 0.083 +/- 0.022 | 0.159 |
| BRIEF | Self-monitor | -0.009 +/- 0.021 | 0.560 |
| BRIEF | Global executive composite | 0.14 +/- 0.018 | 0.036 |
| BSI | Somatization | 0.19 +/- 0.012 | 0.007 |
| BSI | Obsessive-compulsive | 0.097 +/- 0.017 | 0.099 |
| BSI | Interpersonal sensitivity | 0.162 +/- 0.013 | 0.011 |
| BSI | Depression | 0.15 +/- 0.011 | 0.021 |
| BSI | Anxiety | 0.195 +/- 0.010 | 0.003 |
| BSI | Hostility | 0.123 +/- 0.012 | 0.046 |
| BSI | Phobic anxiety | 0.156 +/- 0.014 | 0.011 |
| BSI | Paranoid ideation | 0.141 +/- 0.013 | 0.024 |
| BSI | Psychoticism | 0.134 +/- 0.012 | 0.041 |
| BSI | Global severity index | 0.174 +/- 0.010 | 0.005 |
| BSI | Positive symptom total | 0.158 +/- 0.014 | 0.031 |
| BSI | Positive symptom distress index | 0.078 +/- 0.018 | 0.137 |
| IRI | Fantasy | 0.07 +/- 0.025 | 0.180 |
| IRI | Empathic concern | -0.064 +/- 0.032 | 0.782 |
| IRI | Perspective taking | -0.079 +/- 0.028 | 0.819 |
| IRI | Personal distress | 0.135 +/- 0.021 | 0.034 |
| PANAS | Negative affect | 0.155 +/- 0.016 | 0.020 |
| PANAS | Fear | 0.105 +/- 0.016 | 0.090 |
| PANAS | Sadness | 0.099 +/- 0.015 | 0.087 |
| PANAS | Guilt | 0.142 +/- 0.015 | 0.041 |
| PANAS | Hostility | 0.181 +/- 0.014 | 0.007 |
| PANAS | Shyness | 0.022 +/- 0.025 | 0.412 |
| PANAS | Fatigue | 0.144 +/- 0.015 | 0.031 |
| PANAS | Positive affect | 0.164 +/- 0.014 | 0.019 |
| PANAS | Joviality | 0.184 +/- 0.015 | 0.005 |
| PANAS | Self-assurance | 0.179 +/- 0.014 | 0.007 |
| PANAS | Attentiveness | 0.091 +/- 0.019 | 0.121 |
| PANAS | Serenity | 0.147 +/- 0.017 | 0.027 |
| PANAS | Surprise | 0.147 +/- 0.013 | 0.031 |
| PSQI | Subjective sleep quality | 0.208 +/- 0.012 | 0.005 |
| PSQI | Sleep latency | 0.224 +/- 0.015 | < 0.001 |
| PSQI | Sleep duration | 0.204 +/- 0.009 | < 0.001 |
| PSQI | Habitual sleep efficiency | 0.083 +/- 0.017 | 0.151 |
| PSQI | Sleep disturbance | 0.163 +/- 0.011 | 0.017 |
| PSQI | Use of sleep medication | 0.227 +/- 0.017 | < 0.001 |
| PSQI | Daytime dysfunction | 0.008 +/- 0.026 | 0.478 |
| PSQI | Global sleep quality score | 0.249 +/- 0.012 | < 0.001 |
| PSS | Perceived stress scale | 0.059 +/- 0.022 | 0.235 |

**Supplemental Table 2. Clinical measures' prediction values.** Each clinical measure's prediction strength and *p*-value is shown. Prediction strength is reported as the median Spearman correlation between the observed and predicted measure values across 1000 permutations (+/- 1 standard deviation). *P*-values were generated using permutation testing by comparing the median prediction coefficient to the entire null distribution.

| Assessment | Measure | Cognitive abilities tested | Prediction strength ($\rho$) | P-value (FDR corrected) |
|---|---|---|---|---|
| BNT | Number correct | Lexical retrieval | 0.230 +/- 0.013 | 0.000 |
| BNT | Number of stimulus cues | Lexical retrieval | 0.193 +/- 0.019 | 0.003 |
| BNT | Number correct following stimulus cues | Lexical retrieval | 0.095 +/- 0.021 | 0.121 |
| BNT | Number of phoenomic cues | Lexical retrieval | 0.328 +/- 0.012 | 0.000 |
| BNT | Number correct following phoenomic cues | Lexical retrieval | 0.180 +/- 0.020 | 0.000 |
| BNT | Total score | Lexical retrieval | 0.322 +/- 0.012 | 0.000 |
| WRAT | Word reading | Reading | 0.302 +/- 0.012 | 0.000 |
| WRAML | Verbal learning | Verbal memory | 0.361 +/- 0.008 | 0.000 |
| WRAML | Verbal learning delayed recall | Verbal memory | 0.377 +/- 0.008 | 0.000 |
| WRAML | Verbal learning intrusion | Verbal memory | -0.064 +/- 0.032 | 0.795 |
| WRAML | Finger windows | Visuospatial working memory | 0.380 +/- 0.009 | 0.000 |
| WAIS | Symbol search | Visual perception and processing speed | 0.352 +/- 0.009 | 0.000 |
| WAIS | Coding | Visual-motor and psychomotor speed | 0.397 +/- 0.008 | 0.000 |
| WAIS | Letter-number sequencing | Working memory | 0.302 +/- 0.011 | 0.000 |
| WAIS | Cancellation | Visual processing speed, attention, and visual-motor ability | 0.226 +/- 0.013 | 0.000 |
| D-KEFS | Trail-making test: number sequencing | Processing speed and visual-motor coordination | 0.256 +/- 0.015 | 0.000 |
| D-KEFS | Trail-making test: letter sequencing | Processing speed and visual-motor coordination | 0.346 +/- 0.009 | 0.000 |
| D-KEFS | Trail-making test: number-letter sequencing | Cognitive flexibility and visual-motor coordination | 0.359 +/- 0.010 | 0.000 |
| D-KEFS | Verbal fluency test: letter fluency | Verbal fluency | 0.196 +/- 0.013 | 0.007 |
| D-KEFS | Verbal fluency test: category fluency | Verbal fluency | 0.159 +/- 0.015 | 0.030 |
| D-KEFS | Color-word interference test: color naming | Processing speed | 0.187 +/- 0.015 | 0.005 |
| D-KEFS | Color-word interference test: word reading | Processing speed | 0.192 +/- 0.018 | 0.000 |
| D-KEFS | Color-word interference test: inhibition | Processing speed, cognitive flexibility, inhibition, and switching | 0.302 +/- 0.010 | 0.000 |
| D-KEFS | Twenty-questions test: initial abstraction | Abstract reasoning, concept formation, and problem-solving | 0.374 +/- 0.010 | 0.000 |
| D-KEFS | Twenty-questions test: total questions asked | Abstract reasoning, concept formation, and problem-solving | 0.185 +/- 0.010 | 0.005 |
| D-KEFS | Twenty-questions test: weighted achievement | Abstract reasoning, concept formation, and problem-solving | 0.304 +/- 0.009 | 0.000 |
| WASI | Vocabulary | Vocabulary, concept formation, and expressive language | 0.281 +/- 0.008 | 0.000 |
| WASI | Matrix reasoning | Abstract reasoning, problem-solving , and pattern recognition | 0.428 +/- 0.008 | 0.000 |

**Supplemental Table 3. Cognitive measures' prediction values.** Each cognitive measure's prediction strength associated *p*-value, and a brief description of the cognitive abilities theoretically tested by each measure is shown. Prediction strength is reported as the median Spearman correlation between the observed and predicted measure values across 1000 permutations (+/- 1 standard deviation). *P*-values were generated using permutation testing by comparing the median prediction coefficient to the entire null distribution.

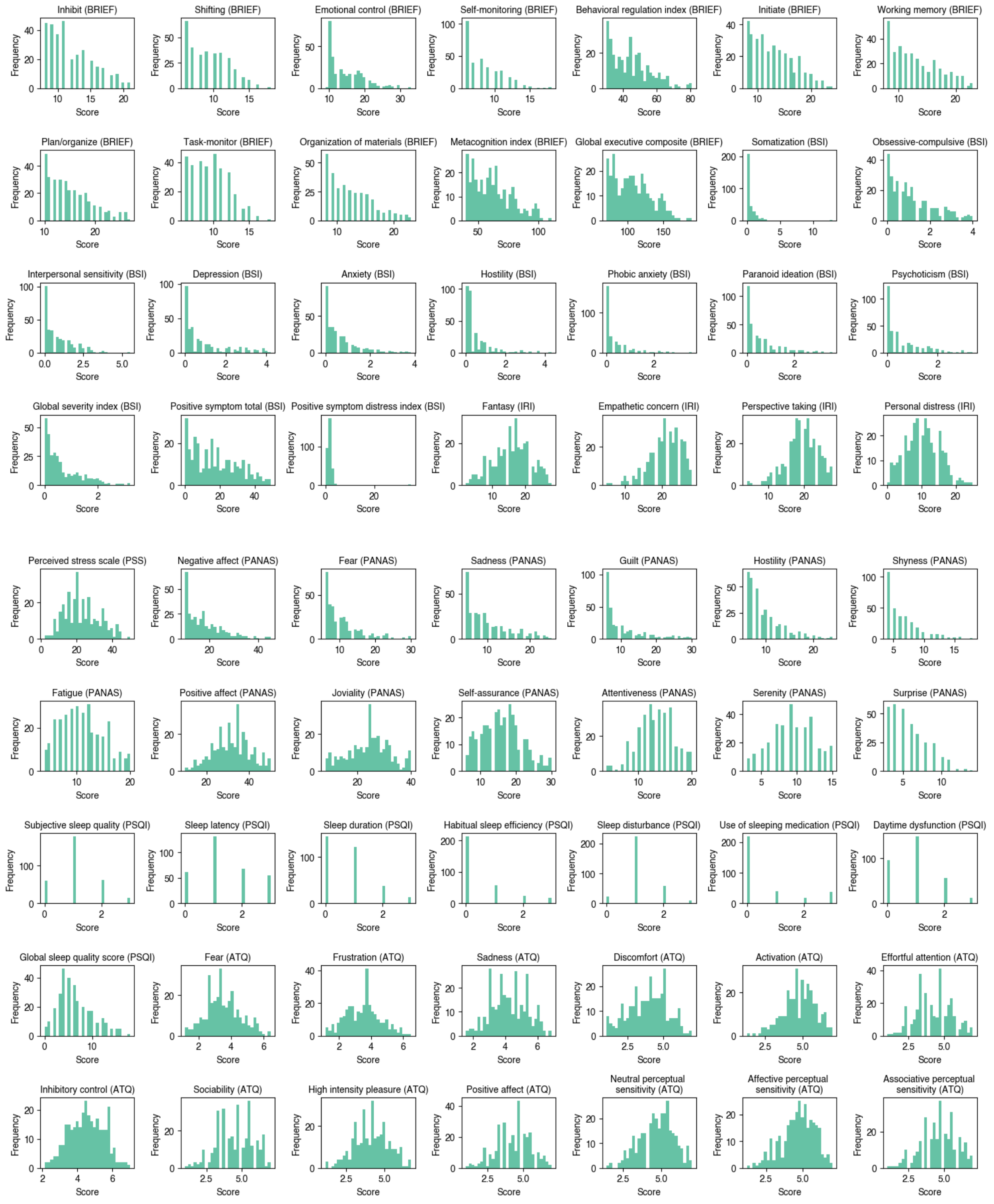


**Supplemental Figure 1. Clinical measures' distributions.** Histograms highlighting the distributions of responses to each clinical measure.

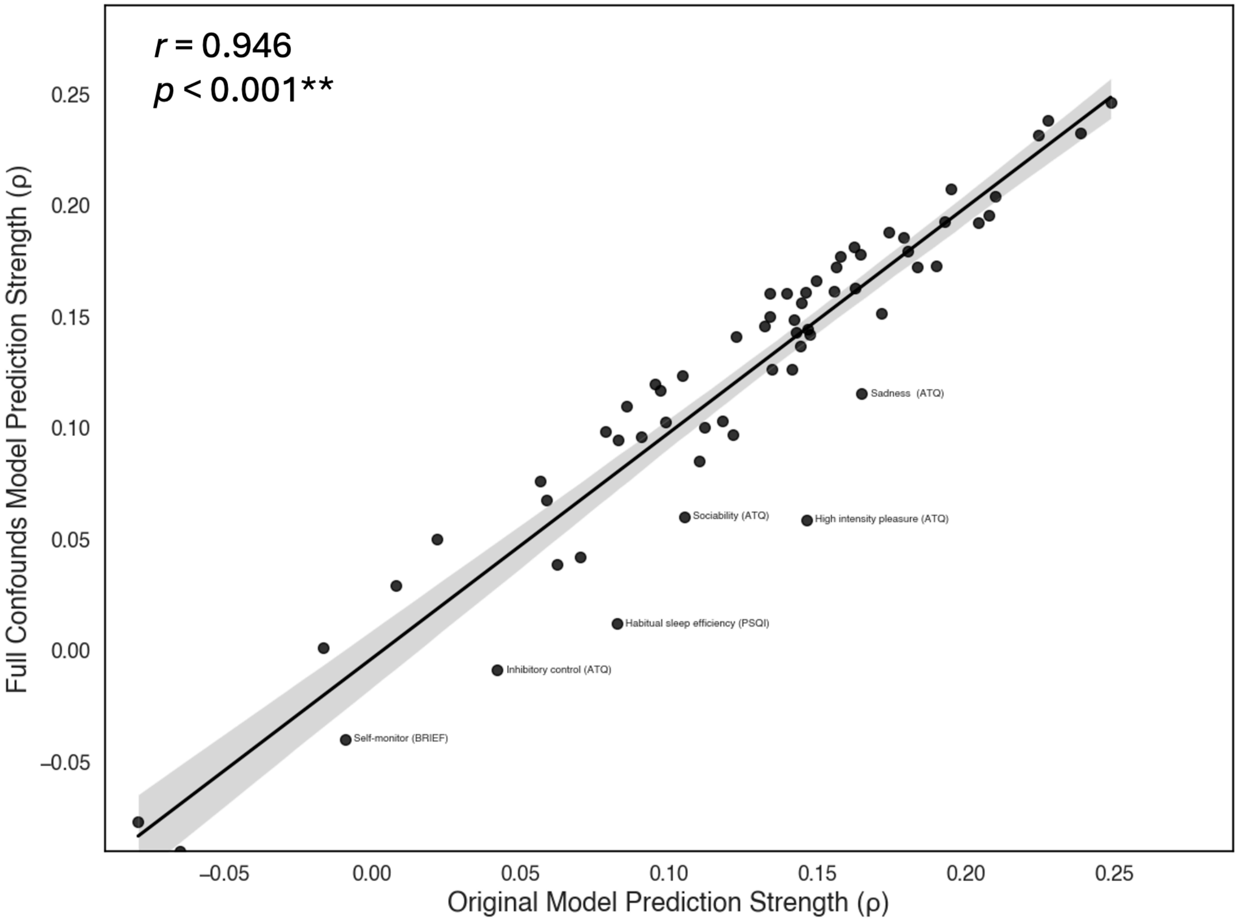


**Supplemental Figure 2. Correlations between prediction strength from the clinical measures' original models compared to prediction strength from models with years of education and income added as covariates.** The correlation between the two models' prediction strengths were high. However, adding education and income covariates in addition to age and sex in the feature selection influenced the prediction strength of several measures. Measures that had either a 0.03 increase or decrease in prediction strength are labeled in the scatterplot.

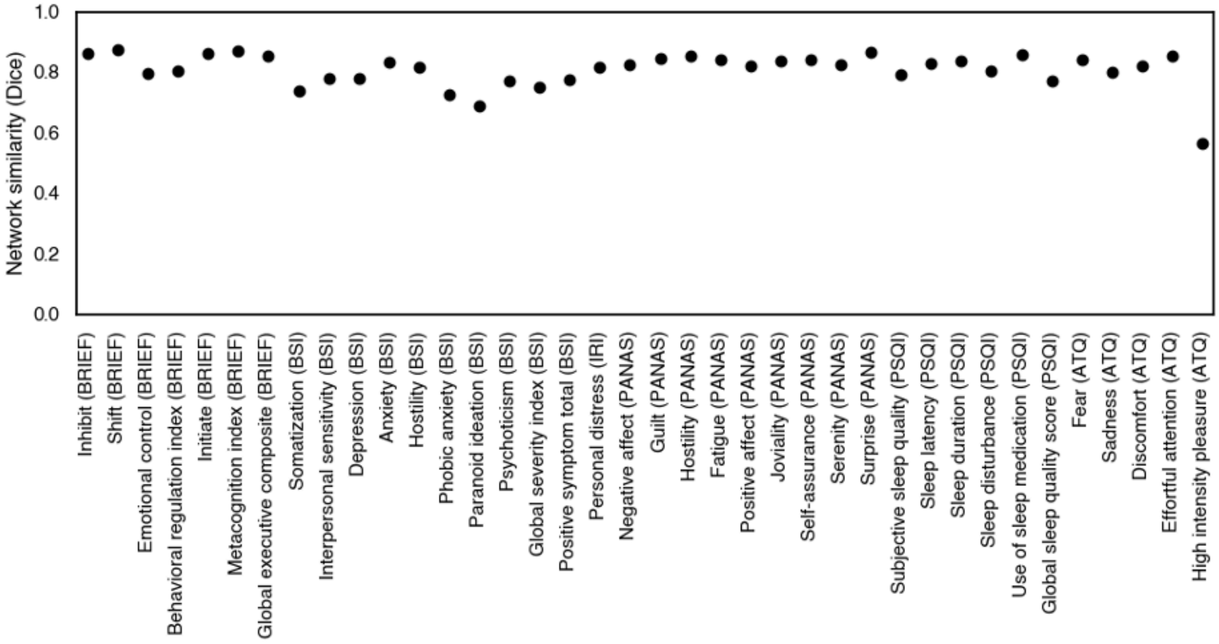


**Supplemental Figure 3. Network similarity between the clinical measures' original models and models with years of education and income added as covariates.** The Dice similarity between the two models' network derivations were typically very high (mean Dice = 0.808 +/- 0.058 stdev).

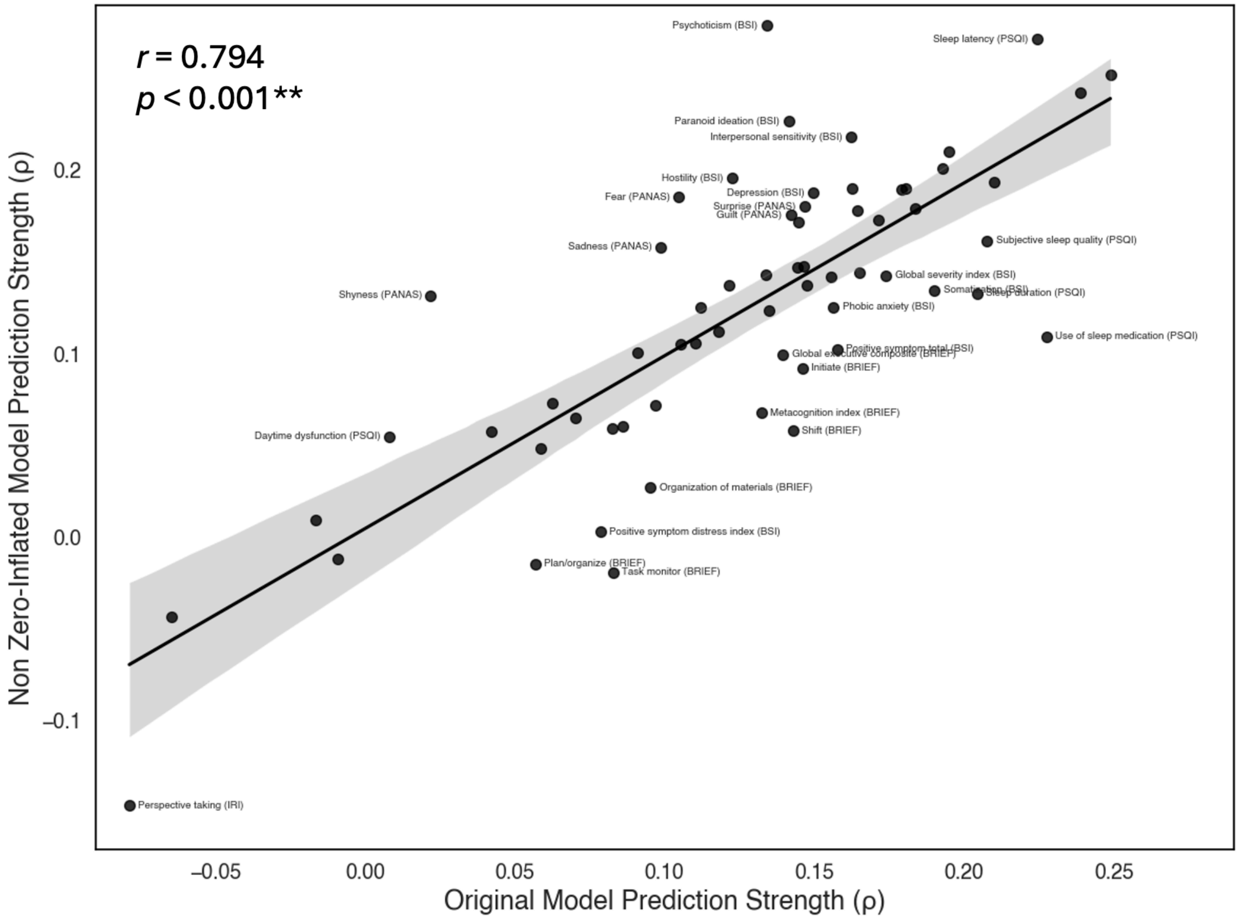


**Supplemental Figure 4. Correlations between clinical measures' prediction strength from original models compared to prediction strength from models re-fit with zero respondents removed.** The correlation between the two models' prediction strengths were high. However, adding education and income covariates in addition to age and sex in the feature selection influenced the prediction strength of several measures. Measures that had either a 0.03 increase or decrease in prediction strength are labeled in the scatterplot.

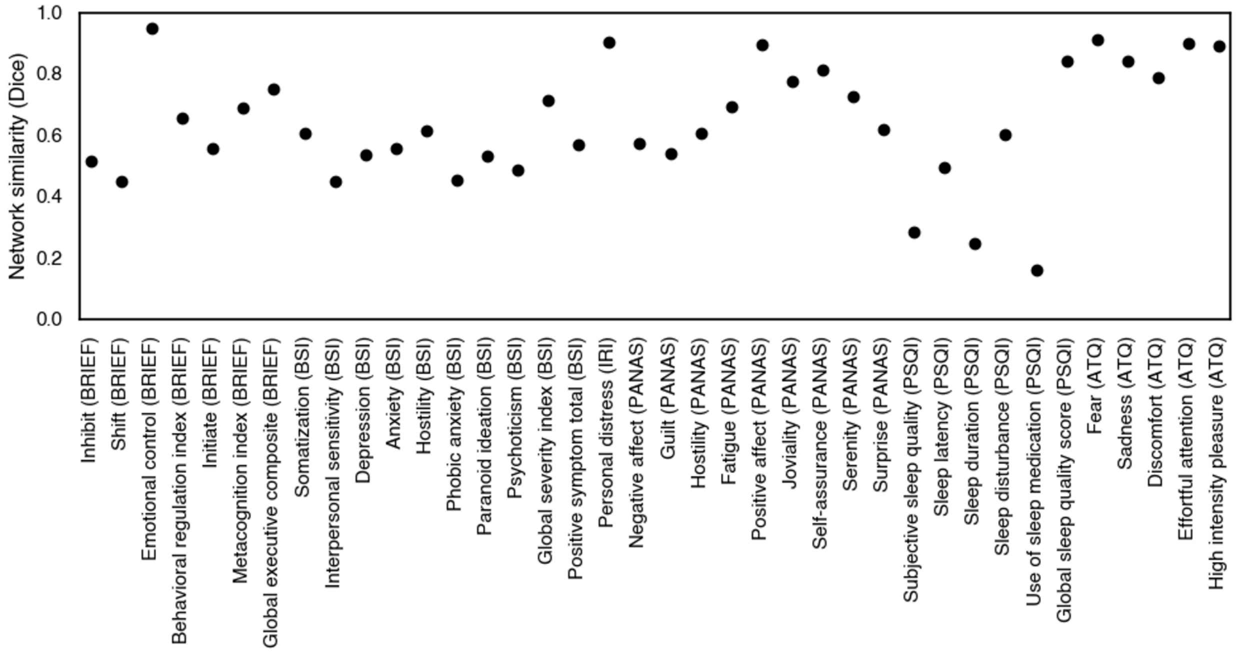


**Supplemental Figure 5. Network similarity between the clinical measures' original models and models re-fit with zero respondents removed.** The Dice similarity between the two models' network derivations were typically high (mean Dice = 0.636 +/- 0.189 stdev). However, networks predictive of subjective sleep quality, sleep duration, and use of sleep medication were less similar (Dice < 0.3).

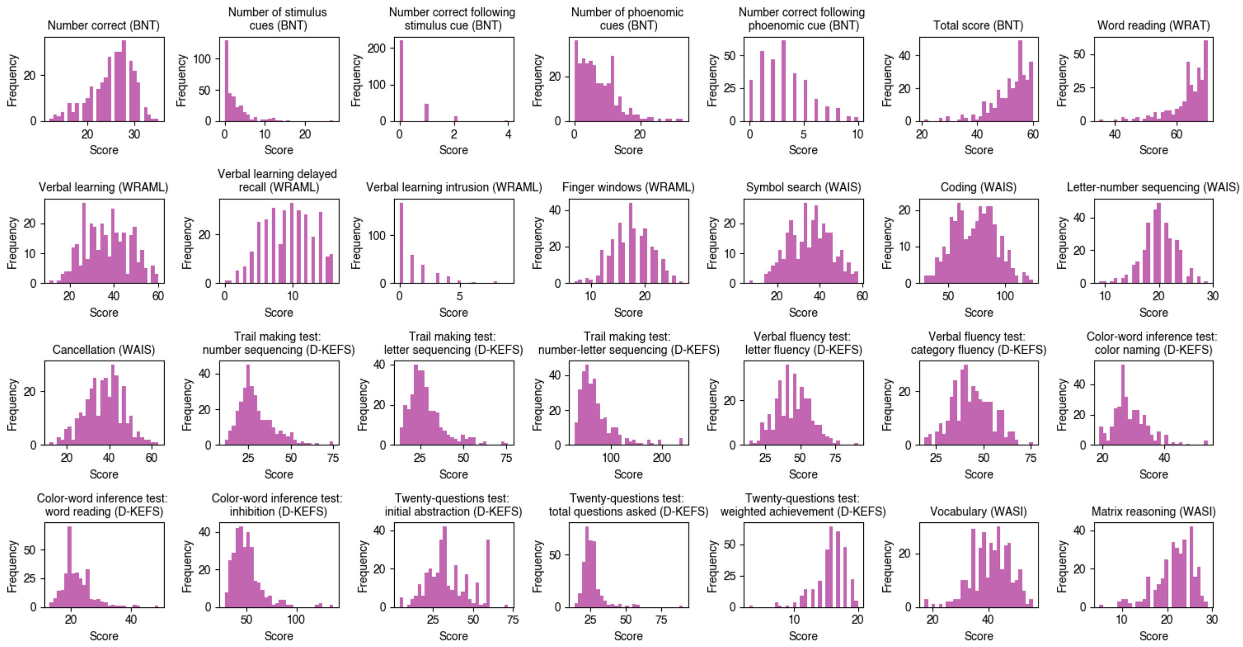


**Supplemental Figure 6. Cognitive measures' distributions.** Histograms highlighting the distributions of scores on each cognitive measure. Note that the two measures with the greatest proportion of scores at ceiling were the two measures the CPM did not significantly predict (number correct following a stimulus cue (BNT) and verbal learning intrusion (WRAML)).

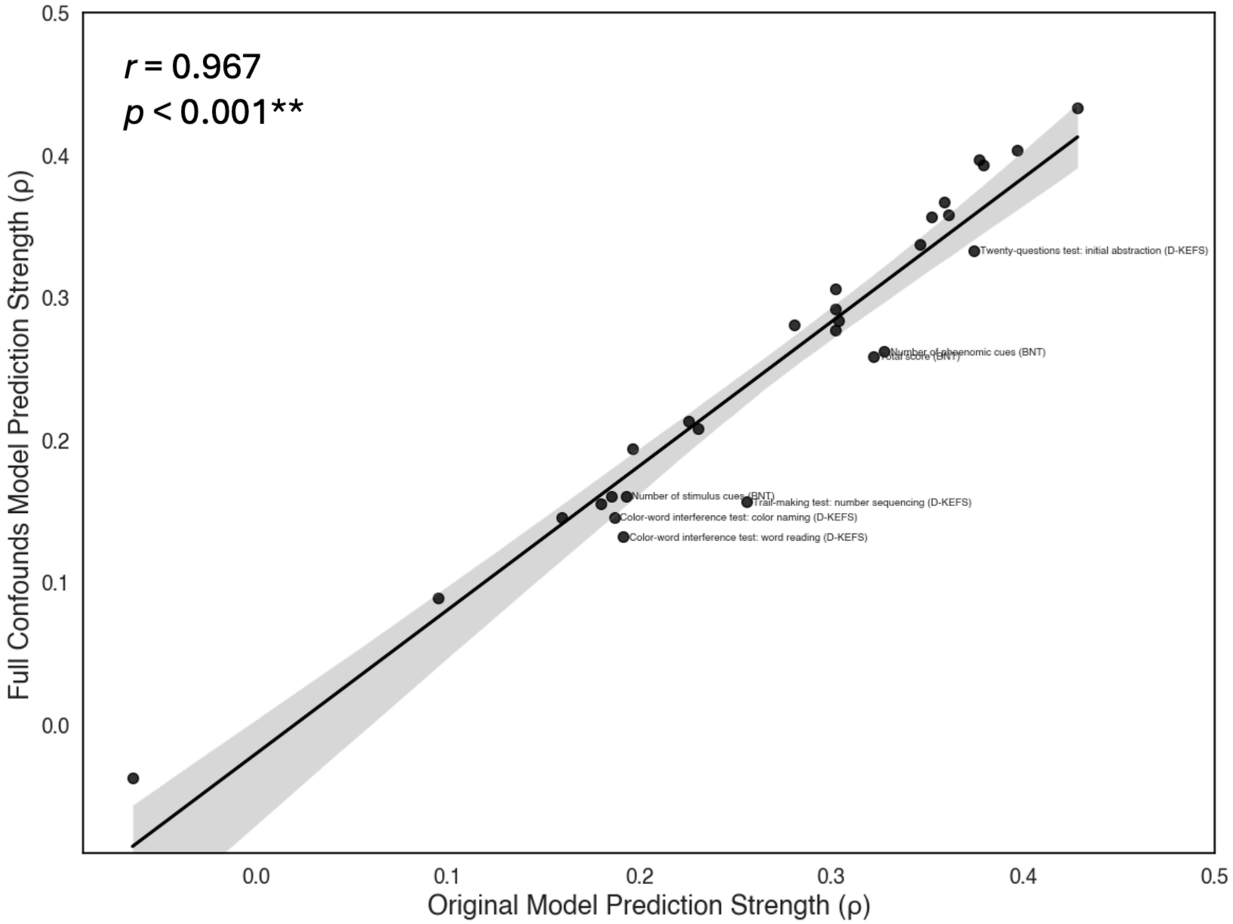


**Supplemental Figure 7. Correlations between prediction strength from the cognitive measures' original models compared to prediction strength from models with years of education and income added as covariates.** The correlation between the two models' prediction strengths were high. However, adding education and income covariates in addition to age and sex in the feature selection influenced the prediction strength of several measures. Measures that had either a 0.03 increase or decrease in prediction strength are labeled in the scatterplot.

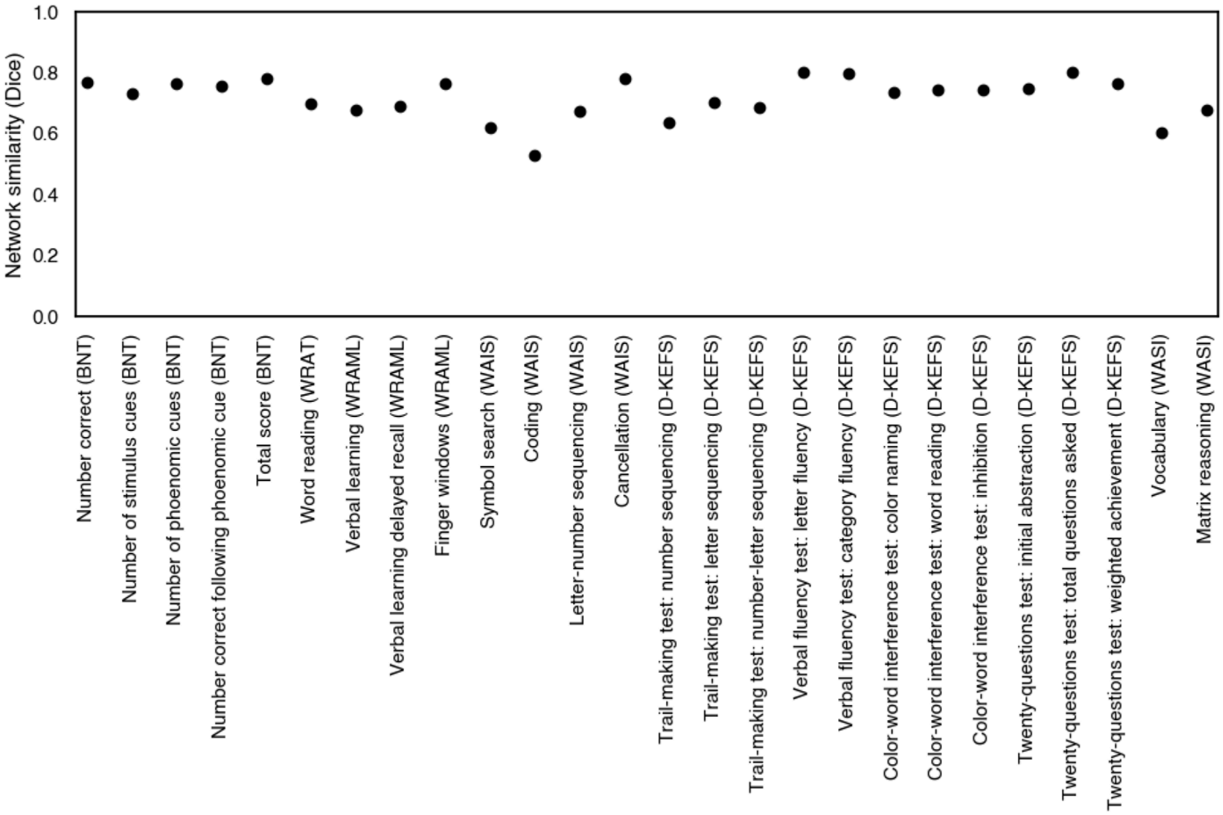


**Supplemental Figure 8. Network similarity between the cognitive measures' original models and models with years of education and income added as covariates.** The Dice similarity between the two models' network derivations were typically high (mean Dice = 0.719 +/- 0.067 stdev).

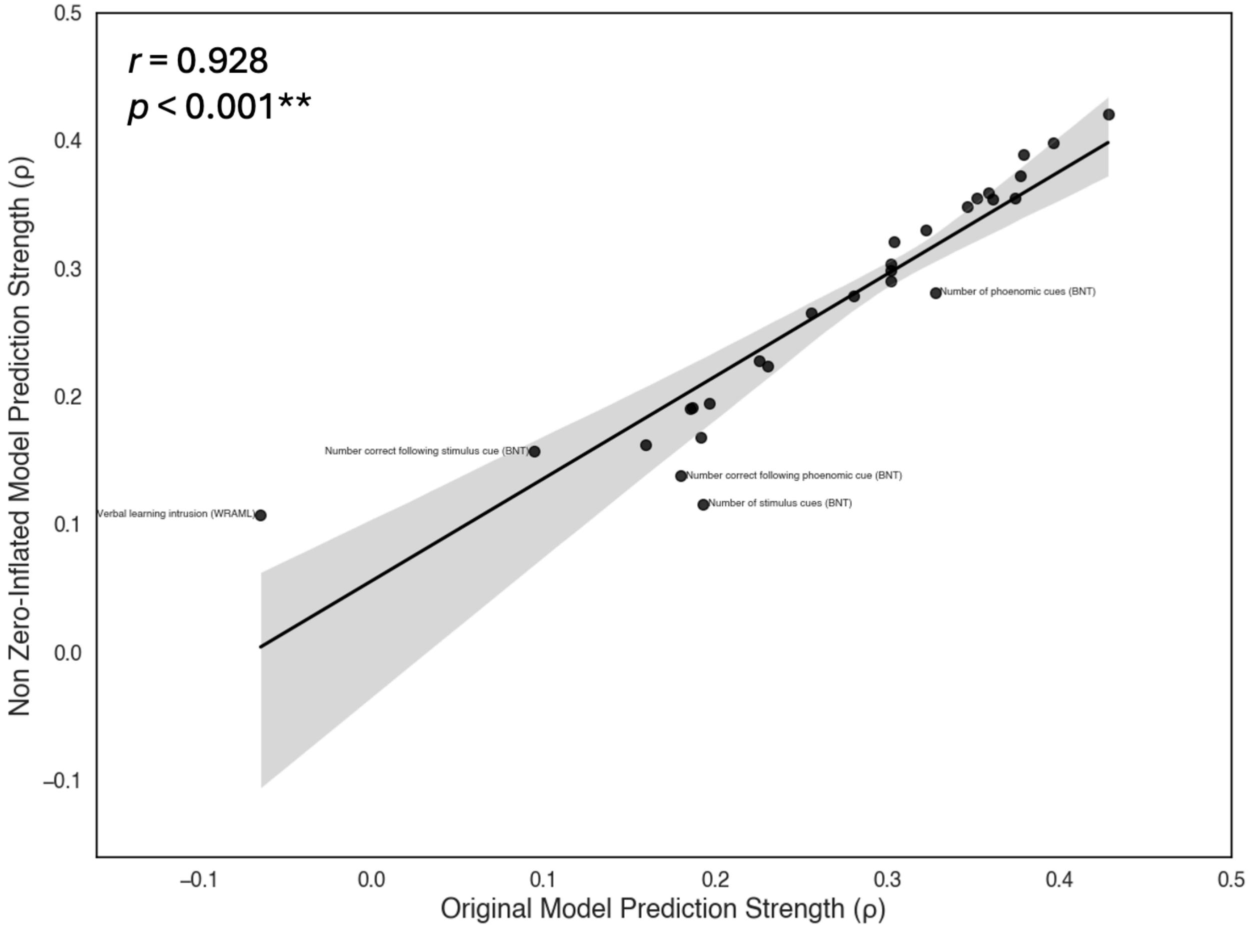


**Supplemental Figure 9. Correlations between cognitive measures' prediction strength from original models compared to prediction strength from models re-fit with zero respondents removed.** The correlation between the two models' prediction strengths were high. However, adding education and income covariates in addition to age and sex in the feature selection influenced the prediction strength of several measures. Measures that had either a 0.03 increase or decrease in prediction strength are labeled in the scatterplot.

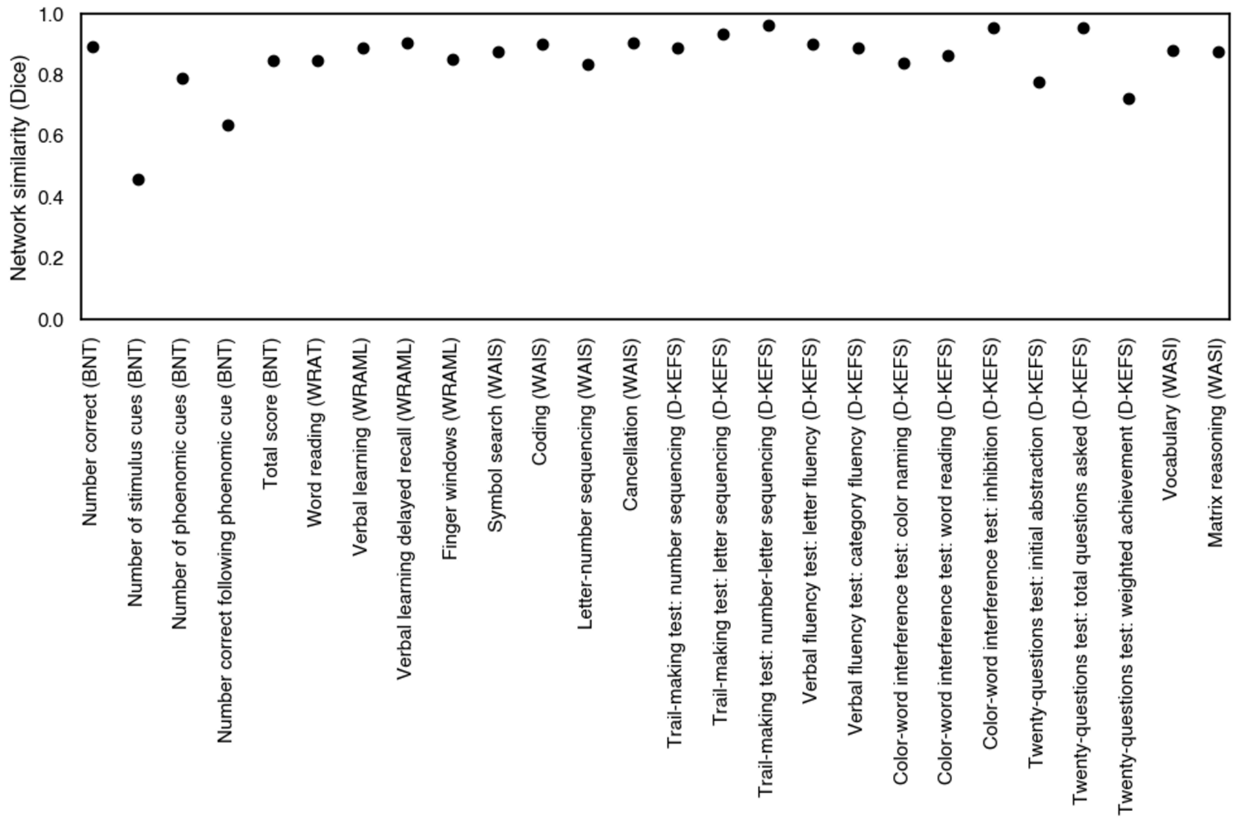


**Supplemental Figure 10. Network similarity between the cognitive measures' original models and models re-fit with zero respondents removed.** The Dice similarity between the two models' network derivations were typically high (mean Dice = 0.849 +/- 0.105 stdev). However, networks predictive of subjective sleep quality, sleep duration, and use of sleep medication were less similar (Dice $< 0.3$).

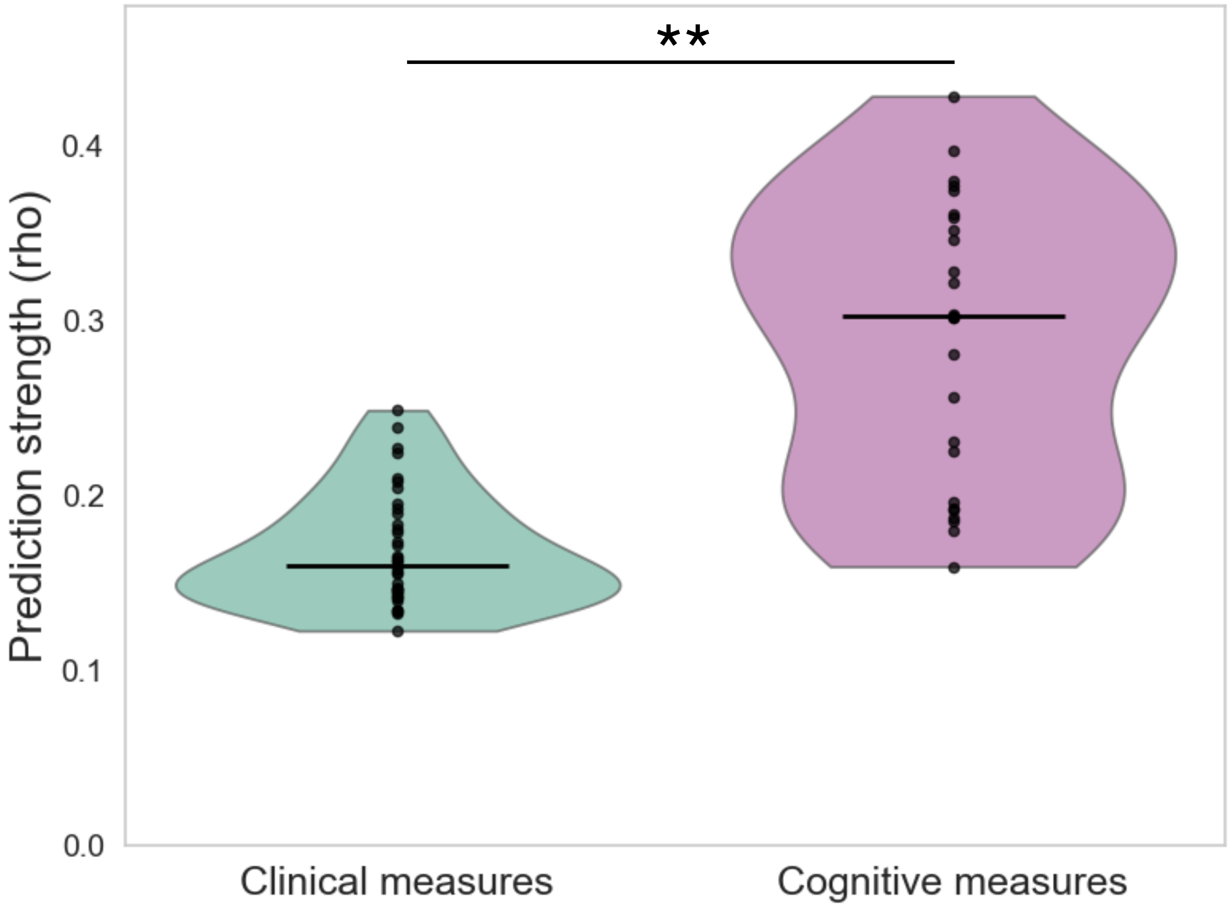


**Supplemental Figure 11. Clinical versus cognitive measure prediction strength.** CPM prediction accuracy of cognitive measures was significantly higher than clinical measure predictions. Horizontal black line = median prediction strength. $^{**}p < 0.001$

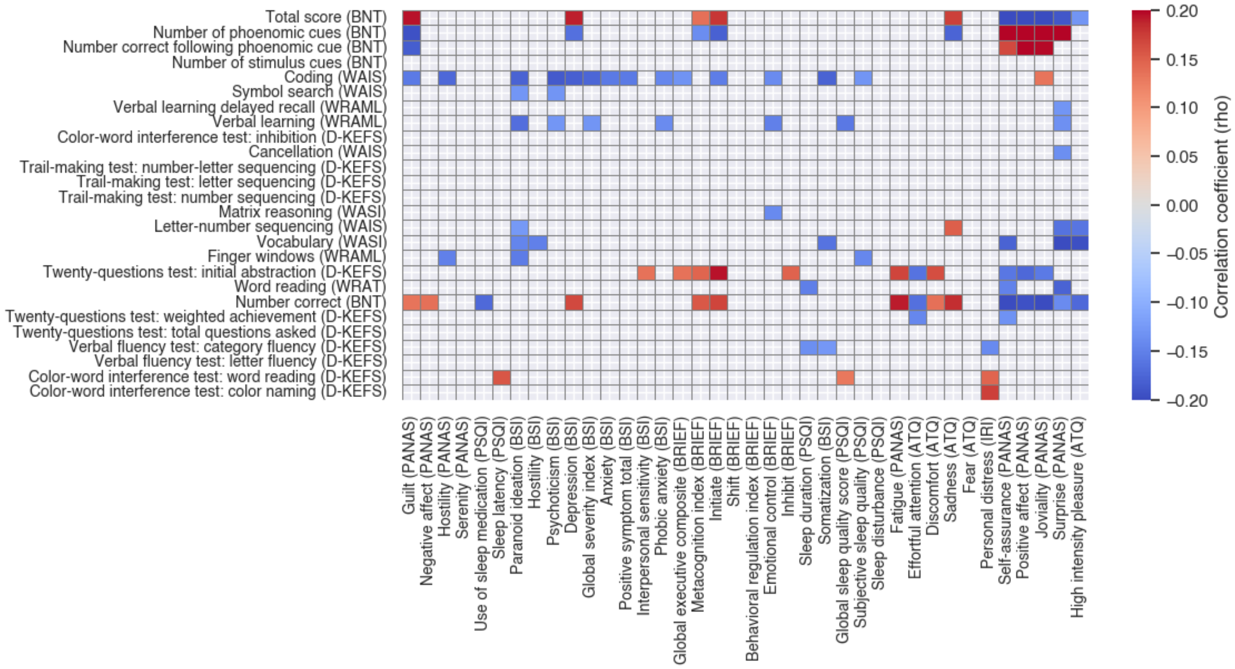


**Supplemental Figure 12. Correlations between clinical and measures**. The Spearman correlation coefficients of the significantly correlated pairs of clinical and cognitive measures are displayed (*pFDR* < 0.05). Of 988 pairwise comparisons, only 101 were significantly correlated. The effect sizes were generally weak (maximum $\rho = 0.234$, minimum $\rho = -0.227$). Measures were sorted according to the network similarity determined by hierarchical clustering.

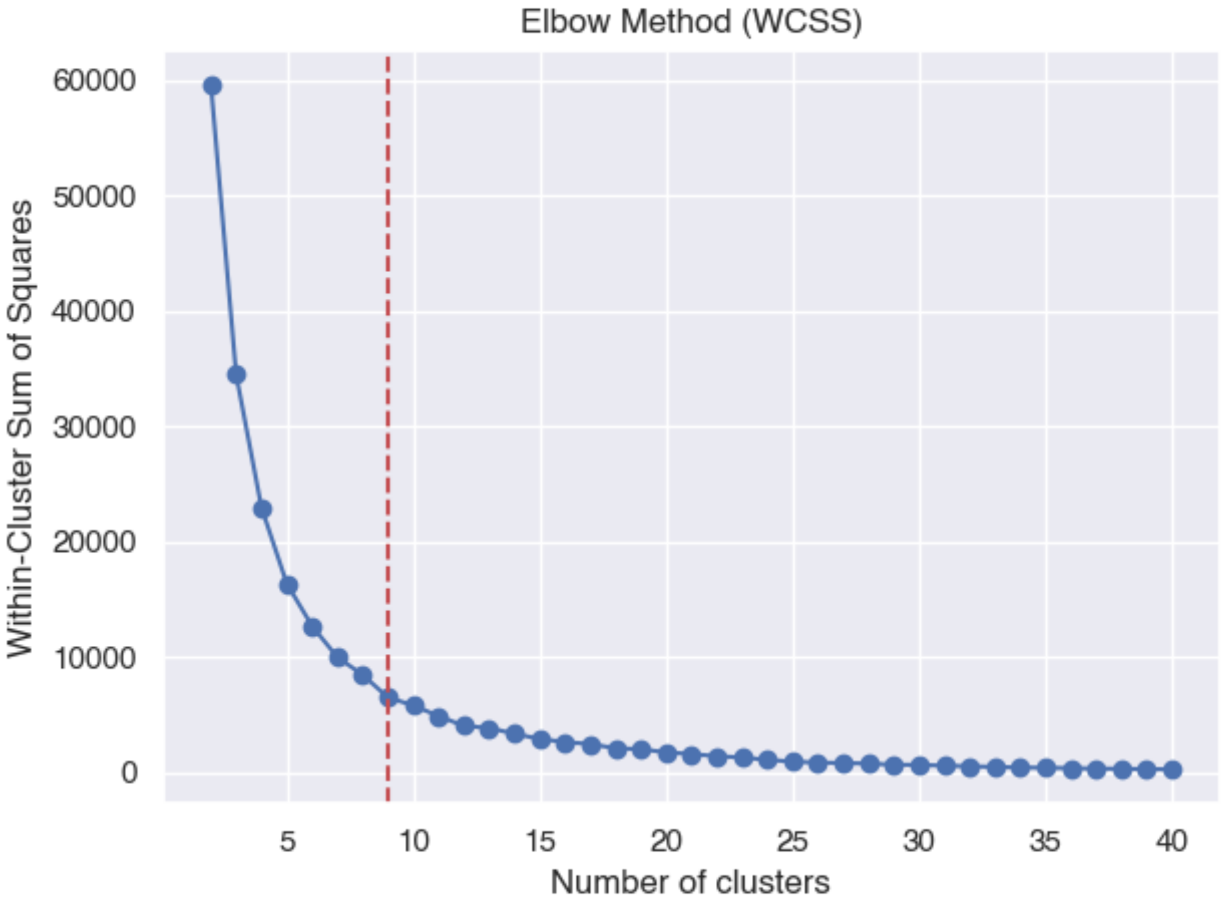


**Supplemental Figure 13. Number of clusters selected using the elbow method.** The curve shows the incremental decrease in within-cluster sum of squares (WCSS) across increasing values of *k* (until 40). The red dashed line denotes the number of clusters (9) selected by identifying the point of maximum curvature (i.e., the “elbow”).

| Psychiatric symptoms | | | Executive functioning | | | Sleep | | |
|---|---|---|---|---|---|---|---|---|
| Shen Atlas Node # | MNI Coordinates | Network | Shen Atlas Node # | MNI Coordinates | Network | Shen Atlas Node # | MNI Coordinates | Network |
| 263 | (-4.87, -10.34, 5.83) | Sub | 113 | (36.7, -57.1, -32.77) | Cer | 12 | (14.33, 36.86, 48.93) | Temp par |
| 128 | (5.46, -9.67, 5.24) | Sub | 107 | (45.96, -46.48, -42.45) | Cer | 6 | (14.56, 64.73, 3.64) | DMN C |
| 40 | (43.34, -10.81, 13.91) | SM B | 102 | (39.09, -74.91, -29.7) | Cer | 217 | (-23.59, -41.29, 19.92) | Sub |
| 253 | (-26.31, -69.52, -30.6) | Cer | 40 | (43.34, -10.81, 13.91) | SM B | 107 | (45.96, -46.48, -42.45) | Cer |
| 172 | (-23.5, -31.62, 63.61) | SM A | 241 | (-40.32, -74.2, -29.15) | Cer | 237 | (-8.7, -50.56, -39.56) | Cer |
| 107 | (45.96, -46.48, -42.45) | Cer | 238 | (-36.95, -52.94, -31.11) | Cer | 14 | (40.68, 14.51, 48.21) | FPCN C |
| 246 | (-42.57, -63.71, -46.29) | Cer | 128 | (5.46, -9.67, 5.24) | Sub | 145 | (-10.15, 55.69, 30.24) | Temp par |
| 159 | (-58.09, -5.56, 27.19) | SM B | 263 | (-4.87, -10.34, 5.83) | Sub | 10 | (8.36, 53.27, 23.91) | DMN C |
| 102 | (39.09, -74.91, -29.7) | Cer | 104 | (23.53, -35.93, -42.96) | Cer | 253 | (-26.31, -69.52, -30.6) | Cer |
| 149 | (-39.35, 17.2, 46.7) | FPCN C | 205 | (-17.04, -50.71, 0.78) | Vis B | 149 | (-39.35, 17.2, 46.7) | FPCN C |
| 23 | (57.78, -8.34, 27.32) | SM B | 149 | (-39.35, 17.2, 46.7) | FPCN C | 141 | (-11.7, 65.09, 4.18) | DMN C |
| 247 | (-10.29, -81.71, -32.29) | Cer | 11 | (37.62, 35.39, 31.09) | VAN B | 194 | (-49.31, -4.7, -37.37) | Temp par |
| 113 | (36.7, -57.1, -32.77) | Cer | 217 | (-23.59, -41.29, 19.92) | Sub | 209 | (-48.34, -67.39, 1.14) | DAN A |
| 33 | (41.97, -23.38, 53.41) | SM A | 133 | (7.47, -34.18, -37.32) | BS | 102 | (39.09, -74.91, -29.7) | Cer |
| 133 | (7.47, -34.18, -37.32) | BS | 267 | (-7.21, -33.04, -39.41) | BS | 33 | (41.97, -23.38, 53.41) | SM A |
| 14 | (40.68, 14.51, 48.21) | FPCN C | 230 | (-32.09, -40.19, -3.99) | Vis B | 40 | (43.34, -10.81, 13.91) | SM B |
| 217 | (-23.59, -41.29, 19.92) | Sub | 253 | (-26.31, -69.52, -30.6) | Cer | 252 | (-46.41, -46.77, -42.86) | Cer |
| 241 | (-40.32, -74.2, -29.15) | Cer | 159 | (-58.09, -5.56, 27.19) | SM B | 263 | (-4.87, -10.34, 5.83) | Sub |
| 230 | (-32.09, -40.19, -3.99) | Vis B | 74 | (45.08, -74.29, 2.58) | Vis A | 128 | (5.46, -9.67, 5.24) | Sub |
| 267 | (-7.21, -33.04, -39.41) | BS | 62 | (39.86, -25.56, 14.38) | SM B | 182 | (-42.05, -65.62, 41.73) | DMN C |
| 104 | (23.53, -35.93, -42.96) | Cer | 180 | (-42.21, -31.23, 15.89) | SM B | 58 | (40.31, -11.27, -35.81) | Lim A |
| 38 | (32.42, -39.19, 49.58) | DAN B | 23 | (57.78, -8.34, 27.32) | SM B | 148 | (-11.17, 34.26, 51.48) | Temp par |
| 114 | (23, -71.84, -29.07) | Cer | 183 | (-51.38, -56.29, 20.47) | Temp par | 68 | (25.23, -44.56, -12.22) | Vis A |
| 99 | (19.29, -7.65, -14.83) | Lim A | 171 | (-50.64, -23.79, 41.37) | DAN B | 11 | (37.62, 35.39, 31.09) | VAN B |
| 158 | (-41.59, -14.68, 44.79) | SM A | 37 | (38.34, -12.45, -1.09) | VAN A | 267 | (-7.21, -33.04, -39.41) | BS |
| 63 | (61.85, -23.77, -2.81) | Temp par | 226 | (-8.79, -42.55, 50.11) | VAN A | 161 | (-6.47, -4.31, 47.6) | SM A |

| Distress | | | Positive affect | | |
|---|---|---|---|---|---|
| Shen Atlas Node # | MNI Coordinates | Network | Shen Atlas Node # | MNI Coordinates | Network |
| 128 | (5.46, -9.67, 5.24) | Sub | 159 | (-58.09, -5.56, 27.19) | SM B |
| 263 | (-4.87, -10.34, 5.83) | Sub | 14 | (40.68, 14.51, 48.21) | FPCN C |
| 238 | (-36.95, -52.94, -31.11) | Cer | 25 | (6.97, -8.07, 52.92) | SM A |
| 113 | (36.7, -57.1, -32.77) | Cer | 23 | (57.78, -8.34, 27.32) | SM B |
| 171 | (-50.64, -23.79, 41.37) | DAN B | 33 | (41.97, -23.38, 53.41) | SM A |
| 159 | (-58.09, -5.56, 27.19) | SM B | 241 | (-40.32, -74.2, -29.15) | Cer |
| 38 | (32.42, -39.19, 49.58) | DAN B | 260 | (-14.6, -3.51, 21.09) | Sub |
| 67 | (36.45, -69.08, -17.46) | Vis A | 100 | (32.22, -78.45, -40.43) | Cer |
| 23 | (57.78, -8.34, 27.32) | SM B | 104 | (23.53, -35.93, -42.96) | Cer |
| 104 | (23.53, -35.93, -42.96) | Cer | 128 | (5.46, -9.67, 5.24) | Sub |
| 237 | (-8.7, -50.56, -39.56) | Cer | 218 | (-7.75, -22.37, 46.05) | SM A |
| 25 | (6.97, -8.07, 52.92) | SM A | 113 | (36.7, -57.1, -32.77) | Cer |
| 132 | (6.31, -24.92, -17.47) | BS | 238 | (-36.95, -52.94, -31.11) | Cer |
| 172 | (-23.5, -31.62, 63.61) | SM A | 184 | (-53.42, -43.53, 38.77) | VAN B |
| 33 | (41.97, -23.38, 53.41) | SM A | 158 | (-41.59, -14.68, 44.79) | SM A |
| 179 | (-35.72, -39.34, 47.75) | DAN B | 24 | (6, -22.28, 65.57) | SM A |
| 265 | (-4.97, -21.51, -15.83) | BS | 239 | (-8.72, -55.18, -52.14) | Cer |
| 253 | (-26.31, -69.52, -30.6) | Cer | 180 | (-42.21, -31.23, 15.89) | SM B |
| 11 | (37.62, 35.39, 31.09) | VAN B | 173 | (-41.23, -15.57, 14.45) | SM B |
| 183 | (-51.38, -56.29, 20.47) | Temp par | 145 | (-10.15, 55.69, 30.24) | Temp par |
| 166 | (-27.58, -9.08, 55.86) | DAN B | 47 | (54.22, -45.24, 36.94) | FPCN C |
| 61 | (59.18, -3.36, 2.74) | SM B | 139 | (-18.21, 56.99, -14.27) | Lim B |
| 74 | (45.08, -74.29, 2.58) | Vis A | 183 | (-51.38, -56.29, 20.47) | Temp par |
| 163 | (-56.98, -3.43, 6.82) | SM B | 122 | (13.75, -4.21, 20.92) | Sub |
| 197 | (-57.05, -14.52, -6.87) | DMN A | 253 | (-26.31, -69.52, -30.6) | Cer |
| 39 | (20.01, -33.24, 69.77) | SM A | 40 | (43.34, -10.81, 13.91) | SM B |

**Supplemental Table 4. Top 10% most important nodes for positively predicting clinical measures.** The most important nodes for positively predicting each clinical measures cluster are shown in descending order, along with the MNI coordinates of the node's centroid and its Yeo 17 networks membership (plus cerebellum, brainstem, and subcortical). 'Cer' = cerebellum, 'BS' = brainstem, 'Sub' = subcortical.

| Psychiatric symptoms | | | Executive functioning | | | Sleep | | |
|---|---|---|---|---|---|---|---|---|
| Shen Atlas Node # | MNI Coordinates | Network | Shen Atlas Node # | MNI Coordinates | Network | Shen Atlas Node # | MNI Coordinates | Network |
| 128 | (5.46, -9.67, 5.24) | Sub | 113 | (36.7, -57.1, -32.77) | Cer | 217 | (-23.59, -41.29, 19.92) | Sub |
| 253 | (-26.31, -69.52, -30.6) | Cer | 241 | (-40.32, -74.2, -29.15) | Cer | 209 | (-48.34, -67.39, 1.14) | DAN A |
| 74 | (45.08, -74.29, 2.58) | Vis A | 102 | (39.09, -74.91, -29.7) | Cer | 167 | (-35.88, -23.3, 65.6) | SM A |
| 263 | (-4.87, -10.34, 5.83) | Sub | 238 | (-36.95, -52.94, -31.11) | Cer | 241 | (-40.32, -74.2, -29.15) | Cer |
| 53 | (52.84, 10.9, -21.83) | DMN A | 253 | (-26.31, -69.52, -30.6) | Cer | 61 | (59.18, -3.36, 2.74) | SM B |
| 40 | (43.34, -10.81, 13.91) | SM B | 107 | (45.96, -46.48, -42.45) | Cer | 84 | (5.25, -1, 35.56) | VAN A |
| 38 | (32.42, -39.19, 49.58) | DAN B | 217 | (-23.59, -41.29, 19.92) | Sub | 237 | (-8.7, -50.56, -39.56) | Cer |
| 113 | (36.7, -57.1, -32.77) | Cer | 128 | (5.46, -9.67, 5.24) | Sub | 204 | (-31.63, -87.16, 12.5) | Vis A |
| 61 | (59.18, -3.36, 2.74) | SM B | 96 | (29.31, -19.59, -26.31) | DMN B | 33 | (41.97, -23.38, 53.41) | SM A |
| 52 | (39.99, 18.91, -34.19) | Lim A | 263 | (-4.87, -10.34, 5.83) | Sub | 120 | (21.22, -36.39, 22.63) | Sub |
| 102 | (39.09, -74.91, -29.7) | Cer | 40 | (43.34, -10.81, 13.91) | SM B | 40 | (43.34, -10.81, 13.91) | SM B |
| 163 | (-56.98, -3.43, 6.82) | SM B | 97 | (24.59, -2.6, -30.72) | Lim A | 163 | (-56.98, -3.43, 6.82) | SM B |
| 114 | (23, -71.84, -29.07) | Cer | 252 | (-46.41, -46.77, -42.86) | Cer | 102 | (39.09, -74.91, -29.7) | Cer |
| 241 | (-40.32, -74.2, -29.15) | Cer | 120 | (21.22, -36.39, 22.63) | Sub | 12 | (14.33, 36.86, 48.93) | Temp par |
| 33 | (41.97, -23.38, 53.41) | SM A | 114 | (23, -71.84, -29.07) | Cer | 172 | (-23.5, -31.62, 63.61) | SM A |
| 247 | (-10.29, -81.71, -32.29) | Cer | 104 | (23.53, -35.93, -42.96) | Cer | 253 | (-26.31, -69.52, -30.6) | Cer |
| 265 | (-4.97, -21.51, -15.83) | BS | 61 | (59.18, -3.36, 2.74) | SM B | 218 | (-7.75, -22.37, 46.05) | SM A |
| 99 | (19.29, -7.65, -14.83) | Lim A | 267 | (-7.21, -33.04, -39.41) | BS | 161 | (-6.47, -4.31, 47.6) | SM A |
| 204 | (-31.63, -87.16, 12.5) | Vis A | 11 | (37.62, 35.39, 31.09) | VAN B | 91 | (8.26, -39.94, 48.11) | FPCN A |
| 97 | (24.59, -2.6, -30.72) | Lim A | 124 | (26.63, 6.34, 0.12) | VAN A | 158 | (-41.59, -14.68, 44.79) | SM A |
| 217 | (-23.59, -41.29, 19.92) | Sub | 133 | (7.47, -34.18, -37.32) | BS | 23 | (57.78, -8.34, 27.32) | SM B |
| 228 | (-26.79, 2.42, -18.71) | Lim A | 14 | (40.68, 14.51, 48.21) | FPCN C | 10 | (8.36, 53.27, 23.91) | DMN C |
| 167 | (-35.88, -23.3, 65.6) | SM A | 149 | (-39.35, 17.2, 46.7) | FPCN C | 146 | (-27.33, 34.07, 36.39) | VAN B |
| 231 | (-22.7, -12.76, -17.43) | DMN B | 12 | (14.33, 36.86, 48.93) | Temp par | 97 | (24.59, -2.6, -30.72) | Lim A |
| 246 | (-42.57, -63.71, -46.29) | Cer | 100 | (32.22, -78.45, -40.43) | Cer | 72 | (20.97, -63.69, -9) | Vis A |
| 92 | (31.16, 3.71, -21.64) | Lim A | 152 | (-28.35, 36.03, -15.64) | Lim B | 145 | (-10.15, 55.69, 30.24) | Temp par |

| Distress | | | Positive affect | | |
|---|---|---|---|---|---|
| Shen Atlas Node # | MNI Coordinates | Network | Shen Atlas Node # | MNI Coordinates | Network |
| 238 | (-36.95, -52.94, -31.11) | Cer | 241 | (-40.32, -74.2, -29.15) | Cer |
| 197 | (-57.05, -14.52, -6.87) | DMN A | 171 | (-50.64, -23.79, 41.37) | DAN B |
| 128 | (5.46, -9.67, 5.24) | Sub | 159 | (-58.09, -5.56, 27.19) | SM B |
| 190 | (-57.62, -6.37, -22.69) | Temp par | 238 | (-36.95, -52.94, -31.11) | Cer |
| 183 | (-51.38, -56.29, 20.47) | Temp par | 33 | (41.97, -23.38, 53.41) | SM A |
| 113 | (36.7, -57.1, -32.77) | Cer | 25 | (6.97, -8.07, 52.92) | SM A |
| 263 | (-4.87, -10.34, 5.83) | Sub | 23 | (57.78, -8.34, 27.32) | SM B |
| 166 | (-27.58, -9.08, 55.86) | DAN B | 267 | (-7.21, -33.04, -39.41) | BS |
| 14 | (40.68, 14.51, 48.21) | FPCN C | 180 | (-42.21, -31.23, 15.89) | SM B |
| 253 | (-26.31, -69.52, -30.6) | Cer | 260 | (-14.6, -3.51, 21.09) | Sub |
| 241 | (-40.32, -74.2, -29.15) | Cer | 133 | (7.47, -34.18, -37.32) | BS |
| 77 | (7.71, -74.97, 25.03) | Vis B | 104 | (23.53, -35.93, -42.96) | Cer |
| 38 | (32.42, -39.19, 49.58) | DAN B | 67 | (36.45, -69.08, -17.46) | Vis A |
| 53 | (52.84, 10.9, -21.83) | DMN A | 167 | (-35.88, -23.3, 65.6) | SM A |
| 52 | (39.99, 18.91, -34.19) | Lim A | 218 | (-7.75, -22.37, 46.05) | SM A |
| 224 | (-7.37, -18.23, 30.02) | FPCN A | 158 | (-41.59, -14.68, 44.79) | SM A |
| 109 | (23.39, -59.27, -52.06) | Cer | 263 | (-4.87, -10.34, 5.83) | Sub |
| 100 | (32.22, -78.45, -40.43) | Cer | 62 | (39.86, -25.56, 14.38) | SM B |
| 186 | (-34.65, 18.63, -32.23) | Lim A | 239 | (-8.72, -55.18, -52.14) | Cer |
| 121 | (12.7, 12.93, 11.49) | Sub | 163 | (-56.98, -3.43, 6.82) | SM B |
| 64 | (56.49, -8.54, -14.27) | DMN C | 128 | (5.46, -9.67, 5.24) | Sub |
| 258 | (-12.52, 11.62, 8.68) | Sub | 38 | (32.42, -39.19, 49.58) | DAN B |
| 252 | (-46.41, -46.77, -42.86) | Cer | 81 | (31.17, -91.77, -10.82) | Vis A |
| 246 | (-42.57, -63.71, -46.29) | Cer | 107 | (45.96, -46.48, -42.45) | Cer |
| 85 | (5.12, -38.94, 27.02) | DMN C | 113 | (36.7, -57.1, -32.77) | Cer |
| 116 | (41.9, -63.98, -49.17) | Cer | 266 | (-4.5, -37.98, -53.13) | BS |

**Supplemental Table 5. Top 10% most important nodes for negatively predicting clinical measures.** The most important nodes for negatively predicting each clinical measures cluster are shown in descending order, along with the MNI coordinates of the node's centroid and its Yeo 17 networks membership (plus cerebellum, brainstem, and subcortical). 'Cer' = cerebellum, 'BS' = brainstem, 'Sub' = subcortical.

| Lexical retrieval | | | Cognitive control | | |
|---|---|---|---|---|---|
| Shen Atlas Node # | MNI Coordinates | Network | Shen Atlas Node # | MNI Coordinates | Network |
| 203 | (-41.25, -75.44, 22.76) | DAN A | 255 | (-6.62, -54.67, -26.32) | Cer |
| 49 | (41.39, -75.34, 27.98) | DAN A | 166 | (-27.58, -9.08, 55.86) | DAN B |
| 113 | (36.7, -57.1, -32.77) | Cer | 25 | (6.97, -8.07, 52.92) | SM A |
| 63 | (61.85, -23.77, -2.81) | Temp par | 254 | (-21.25, -53.41, -23.56) | Cer |
| 100 | (32.22, -78.45, -40.43) | Cer | 110 | (21.07, -54.82, -23.76) | Cer |
| 171 | (-50.64, -23.79, 41.37) | DAN B | 161 | (-6.47, -4.31, 47.6) | SM A |
| 71 | (41.65, -45.73, -22.64) | Vis A | 236 | (-6.52, -66.2, -37.75) | Cer |
| 237 | (-8.7, -50.56, -39.56) | Cer | 199 | (-60.35, -50.04, -14.02) | FPCN C |
| 201 | (-46.67, -39.97, -24.3) | DAN A | 23 | (57.78, -8.34, 27.32) | SM B |
| 239 | (-8.72, -55.18, -52.14) | Cer | 159 | (-58.09, -5.56, 27.19) | SM B |
| 241 | (-40.32, -74.2, -29.15) | Cer | 71 | (41.65, -45.73, -22.64) | Vis A |
| 159 | (-58.09, -5.56, 27.19) | SM B | 34 | (41.79, 4.97, -7.62) | VAN A |
| 197 | (-57.05, -14.52, -6.87) | DMN A | 158 | (-41.59, -14.68, 44.79) | SM A |
| 191 | (-58.98, -29.96, 3.49) | DMN A | 207 | (-25.9, -63.14, -12.25) | Vis A |
| 72 | (20.97, -63.69, -9) | Vis A | 67 | (36.45, -69.08, -17.46) | Vis A |
| 238 | (-36.95, -52.94, -31.11) | Cer | 108 | (16.18, -47.2, -52.31) | Cer |
| 62 | (39.86, -25.56, 14.38) | SM B | 33 | (41.97, -23.38, 53.41) | SM A |
| 104 | (23.53, -35.93, -42.96) | Cer | 243 | (-19.31, -46.44, -53.2) | Cer |
| 33 | (41.97, -23.38, 53.41) | SM A | 72 | (20.97, -63.69, -9) | Vis A |
| 265 | (-4.97, -21.51, -15.83) | BS | 101 | (6.14, -50.74, -12.28) | Cer |
| 264 | (-11.61, -25.62, 14.8) | FPCN A | 59 | (43.36, -26.48, -24.63) | Lim A |
| 78 | (23.7, -96.01, 6.45) | Vis A | 245 | (-27.82, -35.98, -30.88) | Cer |
| 39 | (20.01, -33.24, 69.77) | SM A | 238 | (-36.95, -52.94, -31.11) | Cer |
| 74 | (45.08, -74.29, 2.58) | Vis A | 92 | (31.16, 3.71, -21.64) | Lim A |
| 80 | (7.83, -88.59, 11.86) | Vis A | 89 | (7.83, -23.07, 44.93) | VAN A |
| 32 | (32.05, -5.36, 52.05) | DAN B | 105 | (6.92, -67.95, -37.77) | Cer |

| Fluid intelligence | | | Verbal fluency | | |
|---|---|---|---|---|---|
| Shen Atlas Node # | MNI Coordinates | Network | Shen Atlas Node # | MNI Coordinates | Network |
| 158 | (-41.59, -14.68, 44.79) | SM A | 71 | (41.65, -45.73, -22.64) | Vis A |
| 239 | (-8.72, -55.18, -52.14) | Cer | 192 | (-57.83, -47.48, 5.24) | DMN A |
| 159 | (-58.09, -5.56, 27.19) | SM B | 256 | (-24.3, -37.79, -44.28) | Cer |
| 33 | (41.97, -23.38, 53.41) | SM A | 108 | (16.18, -47.2, -52.31) | Cer |
| 166 | (-27.58, -9.08, 55.86) | DAN B | 23 | (57.78, -8.34, 27.32) | SM B |
| 25 | (6.97, -8.07, 52.92) | SM A | 72 | (20.97, -63.69, -9) | Vis A |
| 171 | (-50.64, -23.79, 41.37) | DAN B | 159 | (-58.09, -5.56, 27.19) | SM B |
| 23 | (57.78, -8.34, 27.32) | SM B | 38 | (32.42, -39.19, 49.58) | DAN B |
| 161 | (-6.47, -4.31, 47.6) | SM A | 212 | (-10.83, -98.14, 7.63) | Vis A |
| 67 | (36.45, -69.08, -17.46) | Vis A | 78 | (23.7, -96.01, 6.45) | Vis A |
| 238 | (-36.95, -52.94, -31.11) | Cer | 110 | (21.07, -54.82, -23.76) | Cer |
| 62 | (39.86, -25.56, 14.38) | SM B | 100 | (32.22, -78.45, -40.43) | Cer |
| 172 | (-23.5, -31.62, 63.61) | SM A | 204 | (-31.63, -87.16, 12.5) | Vis A |
| 191 | (-58.98, -29.96, 3.49) | DMN A | 166 | (-27.58, -9.08, 55.86) | DAN B |
| 237 | (-8.7, -50.56, -39.56) | Cer | 200 | (-42.57, -52.11, -17.36) | DAN A |
| 167 | (-35.88, -23.3, 65.6) | SM A | 260 | (-14.6, -3.51, 21.09) | Sub |
| 61 | (59.18, -3.36, 2.74) | SM B | 119 | (30.36, -36.44, -31.07) | Cer |
| 181 | (-59.51, -25.89, 21.9) | SM B | 114 | (23, -71.84, -29.07) | Cer |
| 110 | (21.07, -54.82, -23.76) | Cer | 105 | (6.92, -67.95, -37.77) | Cer |
| 113 | (36.7, -57.1, -32.77) | Cer | 167 | (-35.88, -23.3, 65.6) | SM A |
| 32 | (32.05, -5.36, 52.05) | DAN B | 73 | (30.1, -82.9, 20.51) | Vis A |
| 199 | (-60.35, -50.04, -14.02) | FPCN C | 187 | (-49.49, 11.11, -30.56) | Temp par |
| 255 | (-6.62, -54.67, -26.32) | Cer | 198 | (-26.69, -42.72, -16.14) | DMN B |
| 71 | (41.65, -45.73, -22.64) | Vis A | 67 | (36.45, -69.08, -17.46) | Vis A |
| 248 | (-8, -68.4, -19.85) | Cer | 206 | (-43.24, -70.37, -13.83) | Vis A |
| 103 | (13.69, -40.06, -25.39) | Cer | 209 | (-48.34, -67.39, 1.14) | DAN A |

**Supplemental Table 6. Top 10% most important nodes for positively predicting cognitive measures.** The most important nodes for positively predicting each cognitive measures cluster are shown in descending order, along with the MNI coordinates of the node's centroid and its Yeo 17 networks membership (plus cerebellum, brainstem, and subcortical). 'Cer' = cerebellum, 'BS' = brainstem, 'Sub' = subcortical.

| Lexical retrieval | | | Cognitive control | | |
|---|---|---|---|---|---|
| Shen Atlas Node # | MNI Coordinates | Network | Shen Atlas Node # | MNI Coordinates | Network |
| 203 | (-41.25, -75.44, 22.76) | DAN A | 63 | (61.85, -23.77, -2.81) | Temp par |
| 49 | (41.39, -75.34, 27.98) | DAN A | 193 | (-59.85, -27.42, -18.14) | FPCN C |
| 63 | (61.85, -23.77, -2.81) | Temp par | 199 | (-60.35, -50.04, -14.02) | FPCN C |
| 145 | (-10.15, 55.69, 30.24) | Temp par | 48 | (47.83, -61.58, 34.71) | DMN C |
| 113 | (36.7, -57.1, -32.77) | Cer | 100 | (32.22, -78.45, -40.43) | Cer |
| 104 | (23.53, -35.93, -42.96) | Cer | 149 | (-39.35, 17.2, 46.7) | FPCN C |
| 191 | (-58.98, -29.96, 3.49) | DMN A | 14 | (40.68, 14.51, 48.21) | FPCN C |
| 201 | (-46.67, -39.97, -24.3) | DAN A | 182 | (-42.05, -65.62, 41.73) | DMN C |
| 197 | (-57.05, -14.52, -6.87) | DMN A | 192 | (-57.83, -47.48, 5.24) | DMN A |
| 62 | (39.86, -25.56, 14.38) | SM B | 191 | (-58.98, -29.96, 3.49) | DMN A |
| 14 | (40.68, 14.51, 48.21) | FPCN C | 143 | (-42.72, 47.25, -6.93) | Temp par |
| 238 | (-36.95, -52.94, -31.11) | Cer | 72 | (20.97, -63.69, -9) | Vis A |
| 239 | (-8.72, -55.18, -52.14) | Cer | 59 | (43.36, -26.48, -24.63) | Lim A |
| 32 | (32.05, -5.36, 52.05) | DAN B | 246 | (-42.57, -63.71, -46.29) | Cer |
| 264 | (-11.61, -25.62, 14.8) | FPCN A | 70 | (60.79, -43.27, -17.64) | FPCN C |
| 237 | (-8.7, -50.56, -39.56) | Cer | 164 | (-23.22, 10.66, 53.61) | FPCN B |
| 147 | (-46.12, 28.15, 26.79) | FPCN B | 55 | (61.28, -22.87, -22.38) | FPCN C |
| 100 | (32.22, -78.45, -40.43) | Cer | 13 | (23.92, 30.67, 36.41) | DMN C |
| 209 | (-48.34, -67.39, 1.14) | DAN A | 114 | (23, -71.84, -29.07) | Cer |
| 13 | (23.92, 30.67, 36.41) | DMN C | 166 | (-27.58, -9.08, 55.86) | DAN B |
| 144 | (-28.81, 50.12, 21.68) | VAN B | 83 | (7.84, 34.68, 17.09) | VAN B |
| 180 | (-42.21, -31.23, 15.89) | SM B | 207 | (-25.9, -63.14, -12.25) | Vis A |
| 234 | (-30.54, -23.92, -26.61) | DMN B | 32 | (32.05, -5.36, 52.05) | DAN B |
| 154 | (-42.97, 42.04, 11.04) | FPCN B | 12 | (14.33, 36.86, 48.93) | Temp par |
| 253 | (-26.31, -69.52, -30.6) | Cer | 36 | (37.5, 21.09, -10.13) | VAN B |
| 33 | (41.97, -23.38, 53.41) | SM A | 22 | (39.98, 17.61, 29.19) | FPCN B |

| Fluid intelligence | | | Verbal fluency | | |
|---|---|---|---|---|---|
| Shen Atlas Node # | MNI Coordinates | Network | Shen Atlas Node # | MNI Coordinates | Network |
| 191 | (-58.98, -29.96, 3.49) | DMN A | 192 | (-57.83, -47.48, 5.24) | DMN A |
| 63 | (61.85, -23.77, -2.81) | Temp par | 190 | (-57.62, -6.37, -22.69) | Temp par |
| 199 | (-60.35, -50.04, -14.02) | FPCN C | 246 | (-42.57, -63.71, -46.29) | Cer |
| 246 | (-42.57, -63.71, -46.29) | Cer | 12 | (14.33, 36.86, 48.93) | Temp par |
| 14 | (40.68, 14.51, 48.21) | FPCN C | 63 | (61.85, -23.77, -2.81) | Temp par |
| 143 | (-42.72, 47.25, -6.93) | Temp par | 72 | (20.97, -63.69, -9) | Vis A |
| 48 | (47.83, -61.58, 34.71) | DMN C | 114 | (23, -71.84, -29.07) | Cer |
| 182 | (-42.05, -65.62, 41.73) | DMN C | 208 | (-16.79, -84.92, 33.04) | Vis B |
| 239 | (-8.72, -55.18, -52.14) | Cer | 149 | (-39.35, 17.2, 46.7) | FPCN C |
| 149 | (-39.35, 17.2, 46.7) | FPCN C | 70 | (60.79, -43.27, -17.64) | FPCN C |
| 193 | (-59.85, -27.42, -18.14) | FPCN C | 187 | (-49.49, 11.11, -30.56) | Temp par |
| 100 | (32.22, -78.45, -40.43) | Cer | 78 | (23.7, -96.01, 6.45) | Vis A |
| 67 | (36.45, -69.08, -17.46) | Vis A | 182 | (-42.05, -65.62, 41.73) | DMN C |
| 166 | (-27.58, -9.08, 55.86) | DAN B | 143 | (-42.72, 47.25, -6.93) | Temp par |
| 238 | (-36.95, -52.94, -31.11) | Cer | 241 | (-40.32, -74.2, -29.15) | Cer |
| 70 | (60.79, -43.27, -17.64) | FPCN C | 193 | (-59.85, -27.42, -18.14) | FPCN C |
| 56 | (54.5, -7.72, -31.53) | Lim A | 204 | (-31.63, -87.16, 12.5) | Vis A |
| 194 | (-49.31, -4.7, -37.37) | Temp par | 102 | (39.09, -74.91, -29.7) | Cer |
| 116 | (41.9, -63.98, -49.17) | Cer | 137 | (-8.15, 39.69, -21.44) | Lim B |
| 253 | (-26.31, -69.52, -30.6) | Cer | 194 | (-49.31, -4.7, -37.37) | Temp par |
| 242 | (-30.17, -80.21, -40.35) | Cer | 242 | (-30.17, -80.21, -40.35) | Cer |
| 113 | (36.7, -57.1, -32.77) | Cer | 42 | (14.82, -68.41, 34.88) | FPCN A |
| 240 | (-21.24, -70.02, -48.88) | Cer | 145 | (-10.15, 55.69, 30.24) | Temp par |
| 71 | (41.65, -45.73, -22.64) | Vis A | 174 | (-7.35, -34.12, 67.46) | SM A |
| 55 | (61.28, -22.87, -22.38) | FPCN C | 199 | (-60.35, -50.04, -14.02) | FPCN C |
| 8 | (44.56, 46.19, -4.9) | FPCN C | 200 | (-42.57, -52.11, -17.36) | DAN A |

**Supplemental Table 7. Top 10% most important nodes for negatively predicting cognitive measures.** The most important nodes for negatively predicting each cognitive measures cluster are shown in descending order, along with the MNI coordinates of the node's centroid and its Yeo 17 networks membership (plus cerebellum, brainstem, and subcortical). 'Cer' = cerebellum, 'BS' = brainstem, 'Sub' = subcortical.

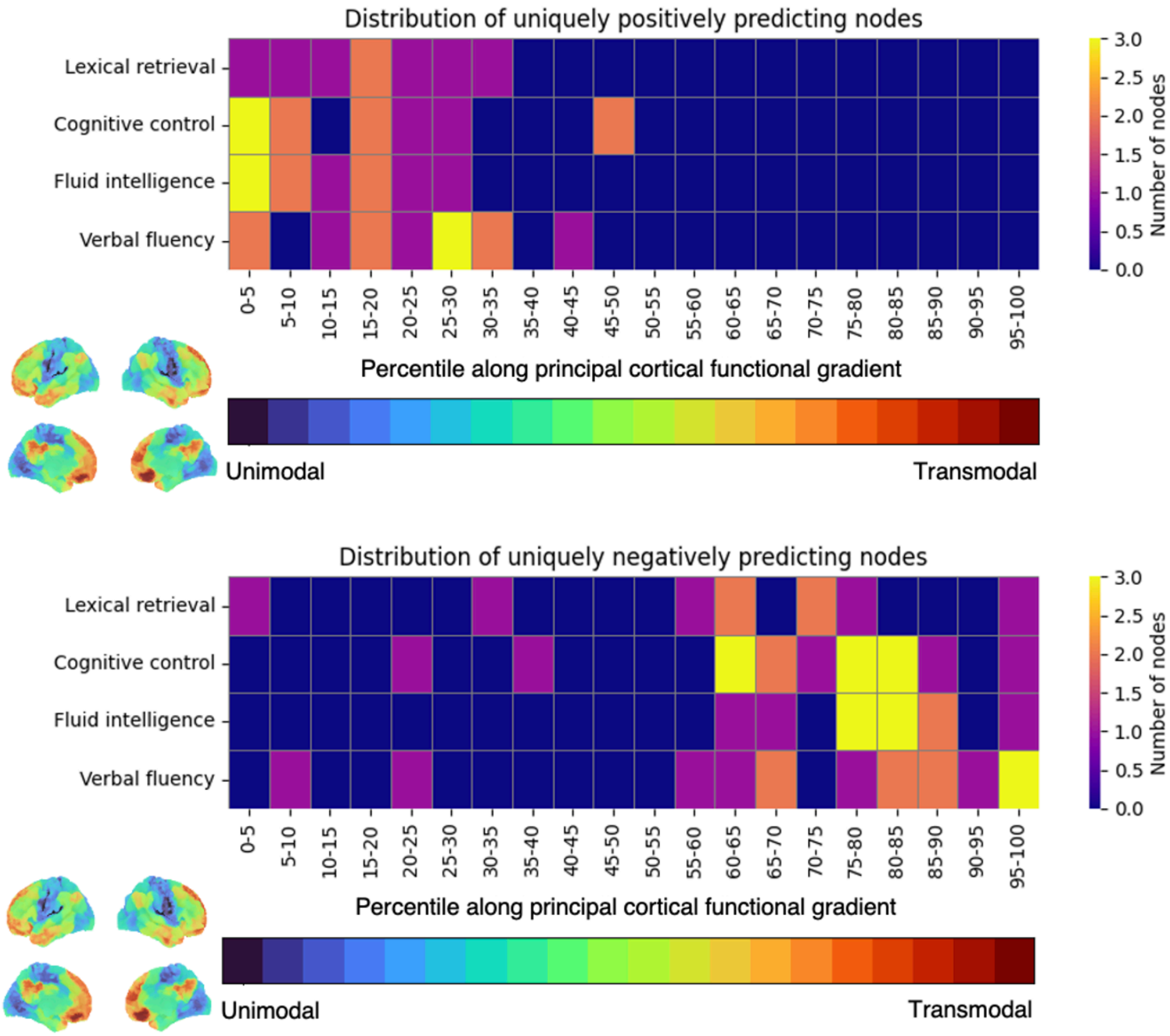


**Supplemental Figure 14. Distribution of nodes important to positively and negatively predicting cognition along the cortical functional hierarchy.** Of the top 10% highest degree predictive nodes for cognition, the ones that were only important to positively predicting cognition were located toward the bottom of the functional gradient in lower-level unimodal cortex, while nodes that were only important in negatively predicting cognition were localized toward the top of the functional gradient in higher-level transmodal cortical regions.

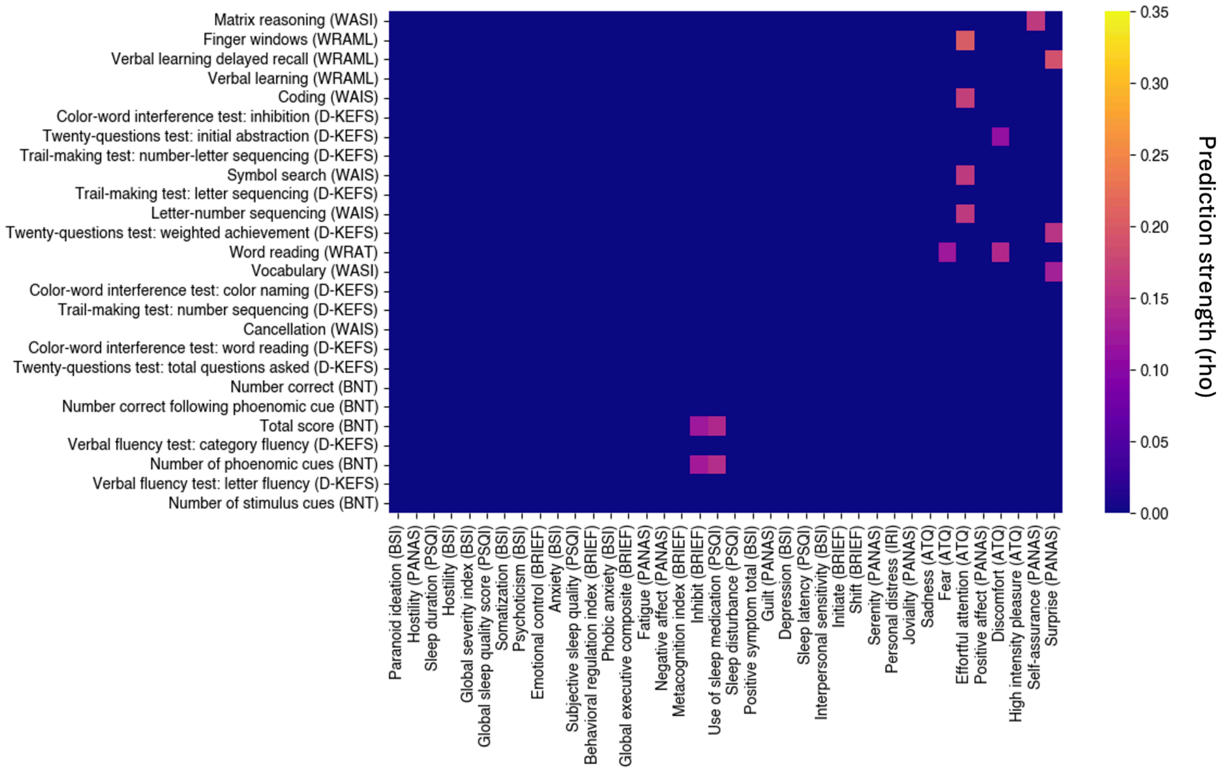


**Supplemental Figure 15. Prediction strength of cognitive measures from randomly selected edges within clinical measure networks.** The prediction strengths were displayed from only the instances where cognitive measure predictions from overlapping edges were not significantly stronger than predictions from a matched number of random edges from clinical networks that did not impinge upon cognitive networks. These instances were confined to sets where overlapping edge predictions were weakest.

| Highest degree nodes from overlapping clinical-cognitive networks that positively predicted cognition | | |
|---|---|---|
| Shen Atlas Node # | MNI Coordinates | Network |
| 149 | (-39.35, 17.2, 46.7) | Temp par |
| 68 | (25.23, -44.56, -12.22) | Vis B |
| 70 | (60.79, -43.27, -17.64) | FPCN B |
| 55 | (61.28, -22.87, -22.38) | FPCN C |
| 193 | (-59.85, -27.42, -18.14) | Temp par |

| Highest degree nodes from overlapping clinical-cognitive networks that negatively predicted cognition | | |
|---|---|---|
| Shen Atlas Node # | MNI Coordinates | Network |
| 194 | (-49.31, -4.7, -37.37) | Lim A |
| 217 | (-23.59, -41.29, 19.92) | Sub |
| 4 | (15.63, 34.11, -22.59) | Lim B |
| 199 | (-60.35, -50.04, -14.02) | FPCN B |
| 120 | (21.22, -36.39, 22.63) | Sub |

**Supplemental Table 8. Top 5 highest degree nodes in overlapping clinical-cognitive networks that predicted cognitive deficits.** The most important nodes to predicting cognition from overlapping clinical-cognitive networks are listed in descending order, along with the MNI coordinates of the node's centroid and its Yeo 17 network membership (plus cerebellum, brainstem, and subcortical). 'Sub' = subcortical.

| Measure | *n* |
|---|---|
| Inhibit (BRIEF) | 316 |
| Shift (BRIEF) | 317 |
| Emotional control (BRIEF) | 316 |
| Self-monitor (BRIEF) | 317 |
| Behavioral regulation index (BRIEF) | 316 |
| Initiate (BRIEF) | 317 |
| Working memory (BRIEF) | 317 |
| Plan/organize (BRIEF) | 317 |
| Task monitor (BRIEF) | 315 |
| Organization of materials (BRIEF) | 317 |
| Metacognition index (BRIEF) | 315 |
| Global executive composite (BRIEF) | 313 |
| Somatization (BSI) | 316 |
| Obsessive-compulsive (BSI) | 316 |
| Interpersonal sensitivity (BSI) | 316 |
| Depression (BSI) | 316 |
| Anxiety (BSI) | 315 |
| Hostility (BSI) | 316 |
| Phobic anxiety (BSI) | 316 |
| Paranoid ideation (BSI) | 316 |
| Psychoticism (BSI) | 316 |
| Global severity index (BSI) | 316 |
| Positive symptom total (BSI) | 316 |
| Positive symptom distress index (BSI) | 315 |
| Fantasy (IRI) | 316 |
| Empathic concern (IRI) | 313 |
| Perspective taking (IRI) | 316 |
| Personal distress (IRI) | 314 |
| Perceived stress scale (PSS) | 313 |

| Measure | *n* |
|---|---|
| Negative affect (PANAS) | 317 |
| Fear (PANAS) | 317 |
| Sadness (PANAS) | 317 |
| Guilt (PANAS) | 317 |
| Hostility (PANAS) | 314 |
| Shyness (PANAS) | 316 |
| Fatigue (PANAS) | 317 |
| Positive affect (PANAS) | 316 |
| Joviality (PANAS) | 317 |
| Self-assurance (PANAS) | 317 |
| Attentiveness (PANAS) | 316 |
| Serenity (PANAS) | 317 |
| Surprise (PANAS) | 317 |
| Subjective sleep quality (PSQI) | 314 |
| Sleep latency (PSQI) | 316 |
| Sleep duration (PSQI) | 317 |
| Habitual sleep efficiency (PSQI) | 314 |
| Sleep disturbance (PSQI) | 316 |
| Use of sleep medication (PSQI) | 313 |
| Daytime dysfunction (PSQI) | 314 |
| Global sleep quality score (PSQI) | 309 |
| Fear (ATQ) | 306 |
| Frustration (ATQ) | 306 |
| Sadness (ATQ) | 306 |
| Discomfort (ATQ) | 306 |
| Activation control (ATQ) | 306 |
| Effortful attention (ATQ) | 306 |
| Inhibitory control (ATQ) | 306 |
| Sociability (ATQ) | 306 |
| High intensity pleasure (ATQ) | 306 |
| Positive affect (ATQ) | 306 |
| Neutral perceptual sensitivity (ATQ) | 306 |
| Affective perceptual sensitivity (ATQ) | 306 |
| Associative perceptual sensitivity (ATQ) | 306 |

**Supplemental Table 9. Number of subjects who had data for each clinical measure.**

| **Measure** | ***n*** |
|---|---|
| Number correct (BNT) | 312 |
| Number of stimulus cues (BNT) | 312 |
| Number correct following stimulus cue (BNT) | 285 |
| Number of phoenomic cues (BNT) | 312 |
| Number correct following phoenomic cue (BNT) | 304 |
| Total score (BNT) | 312 |
| Word reading (WRAT) | 317 |
| Verbal learning (WRAML) | 317 |
| Verbal learning delayed recall (WRAML) | 317 |
| Verbal learning intrusion (WRAML) | 316 |
| Finger windows (WRAML) | 317 |
| Symbol search (WAIS) | 316 |
| Coding (WAIS) | 316 |
| Letter-number sequencing (WAIS) | 317 |
| Cancellation (WAIS) | 317 |
| Trail-making test: number sequencing (D-KEFS) | 317 |
| Trail-making test: letter sequencing (D-KEFS) | 317 |
| Trail-making test: number-letter sequencing (D-KEFS) | 316 |
| Verbal fluency test: letter fluency (D-KEFS) | 315 |
| Verbal fluency test: category fluency (D-KEFS) | 312 |
| Color-word interference test: color naming (D-KEFS) | 315 |
| Color-word interference test: word reading (D-KEFS) | 315 |
| Color-word interference test: inhibition (D-KEFS) | 315 |
| Twenty-questions test: initial abstraction (D-KEFS) | 314 |
| Twenty-questions test: total questions asked (D-KEFS) | 314 |
| Twenty-questions test: weighted achievement (D-KEFS) | 314 |
| Vocabulary (WASI) | 317 |
| Matrix reasoning (WASI) | 317 |

**Supplemental Table 10. Number of subjects who had data for each cognitive measure.**
